# A complex balance: PLETHORA - JANUS ratiometry explains the PLETHORA dosage regeneration paradox

**DOI:** 10.64898/2026.10.15.759210

**Authors:** Monica L Garcia-Gomez, Mabel Maria Mathew, Srijan Das, Chris Roelofsen, Viola Willemsen, Kalika Prasad, Kirsten ten Tusscher

## Abstract

Molecular gradients are classically thought to encode positional information through concentration thresholds, but how transcription factor dosage is translated into regenerative competence remains unclear. In Arabidopsis roots, PLETHORA (PLT) transcription factors form a gradient that defines a regeneration competence zone and, following injury, rapidly re-establishes their transcriptional domain through an auto-activation. Yet, increasing PLT levels has a paradoxical effect: moderate increases expand the competence zone, whereas further increases impair regeneration. Here, combining mathematical modelling with experiments, we uncover the molecular basis of this non-monotonic response. Our model reveals that PLT activity depends not simply on its abundance, but on the balance between PLT protein complexes and unbound PLT. Following injury, PLT auto-activation rapidly restores the transcriptional domain, whereas increasing PLT dosage progressively shifts the equilibrium towards unbound PLT, reducing the availability of active PLT complexes. We identify and experimentally validate a strategy to extend regeneration competence by maintaining an appropriate ratio of PLT complexes to unbound PLT. Our findings reveal a ratiometric mechanism for decoding transcription factor dosage, in which excess regulator can attenuate rather than enhance activity. Such molecular balancing provides a mechanism by which a single transcription factor gradient can generate distinct, context-dependent outcomes during development and regeneration.

## Introduction

Organ regeneration is a remarkable process in which cells near a wound respond by growing, dividing, and changing their identity to rebuild the missing tissues in the correct spatial arrangement. In many model organisms, regeneration relies on the ability of differentiated cells to reprogram and regain developmental potential (Birnbaum and Alvarado, 2008; Mathew and Prasad, 2021). This cell fate flexibility is essential for successful regeneration after damage. Regeneration begins with the acquisition of regeneration competence followed by tissue repatterning process until the original shape and function of the organ are restored (Birnbaum and Alvarado, 2008; Efroni et al., 2016; Mathew and Prasad, 2021; Omary et al., 2023; Mathew et al., 2025).

The Arabidopsis root is an excellent model for studying the acquisition of regeneration competence because different portions of the root tip can be removed with high precision, enabling the spatio-temporal analysis of the remaining cells and their regenerative responses (Sena et al., 2009; Xu et al., 2006; Efroni et al., 2016; Kral et al., 2016; Heyman et al., 2016; Durgaprasad et al., 2019; Matosevich et al., 2020; Fallesen et al., 2024; Lee et al., 2025; Rahni et al., 2026; Garcia-Gomez and Ten Tusscher, 2024; Baulies et al., 2025). The root apex is developmentally organized along its longitudinal axis. Stem cells at the tip continuously generate daughter cells that divide rapidly within the meristem, undergo anisotropic expansion in the elongation zone, and ultimately differentiate in the differentiation zone. Molecular gradients provide positional information that guides root cells through these developmental zones, with quantitative differences in molecular signals contributing to their spatial organization and progressive changes in cell fate. Auxin and the PLETHORA (PLT) transcription factors both form gradients along the root apex, reaching their maximum levels in the root stem cell niche (SCN). Whereas PLT dosage transcription-factors regulate cell identity, stem cell maintenance, cell division and elongation, thereby establishing the zonation of the root apex, auxin levels primarily modulate the rates of these processes (Aida et al., 2004; Galinha et al., 2007; Mähönen et al., 2014; Santuari et al., 2016).

Removal of the distal end of the root tip, from the columella to a few cells above the stem cell niche (SCN) (i.e. low cuts, LC), triggers a series of wound responses that regenerate all the missing cell types, and restores the characteristic convex root tip within 2–3 days (Xu et al., 2006; Efroni et al., 2016; Mathew et al., 2025). Notably, competence to regenerate declines as the cut is made further away from the SCN (i.e. high cuts, HC) (Sena et al., 2009; Durgaprasad et al., 2019). This positional effect defines a regeneration competence zone, and suggests that regeneration depends on molecular factors that are enriched near the root tip. Consistently, the plant hormone auxin and the auxin-regulated PLT transcription factor have emerged as key regulators of regeneration competence (Durgaprasad et al., 2019; Matosevich et al., 2020). Either exogenous auxin or a short (2 hours) induction of PLT2 nuclear localization before root tip cutting enables successful regeneration after HC that would otherwise fail (Durgaprasad et al., 2019; Matosevich et al., 2020). Similarly, *MIR396* overexpression, that elevates PLT levels (Rodriguez et al., 2015), also enhances regeneration competence (Baulies et al., 2025). Rather than acting independently, we hypothesise that the effects of auxin and PLT on regeneration competence can be explained by the positive feedback loop between them (Santuari et al., 2016; Salvi et al., 2020), such that they function as a single integrated regulatory module controlling regeneration. Paradoxically, prolonged induction of PLT2 nuclear localisation (48 hours) before cutting reverses the beneficial effect observed after a short induction, resulting in reduced regeneration competence at HC (Durgaprasad et al., 2019). The molecular basis of this time- and dosage-dependent effect remains unclear and suggests that the auxin–PLT regulatory network is more complex than currently understood. This interpretation is consistent with the recent identification of multiple regulatory regions within the *PLT* promoter (Kerstens et al., 2025), suggesting multi-layer regulation of PLT expression.

Here we combine computational modelling with experiments to resolve the paradoxical effects of PLT induction on root regeneration competence. Our model incorporates PLT protein complexes that control PLT auto-activation and the expression of downstream genes (Figure 1). PLT auto-activation is mediated by the PLT-JANUS complex. We show that prolonged PLT induction progressively shifts the balance between these molecular states, increasing unbound PLT and thereby limiting the activity of transcriptionally active PLT-JANUS complexes. This provides a mechanistic explanation for why increasing PLT levels can initially enhance, but ultimately impair, regeneration competence. Guided by this model, we identify and experimentally validate a strategy to extend regeneration competence by maintaining an appropriate balance between PLT protein complexes and unbound PLT. Together, our findings establish ratiometric control of PLT activity as a mechanism for decoding transcription factor dosage, enabling PLT to promote regeneration over short timescales while maintaining robust control of developmental transitions and long-term stem cell behaviour.

**Figure 1.**
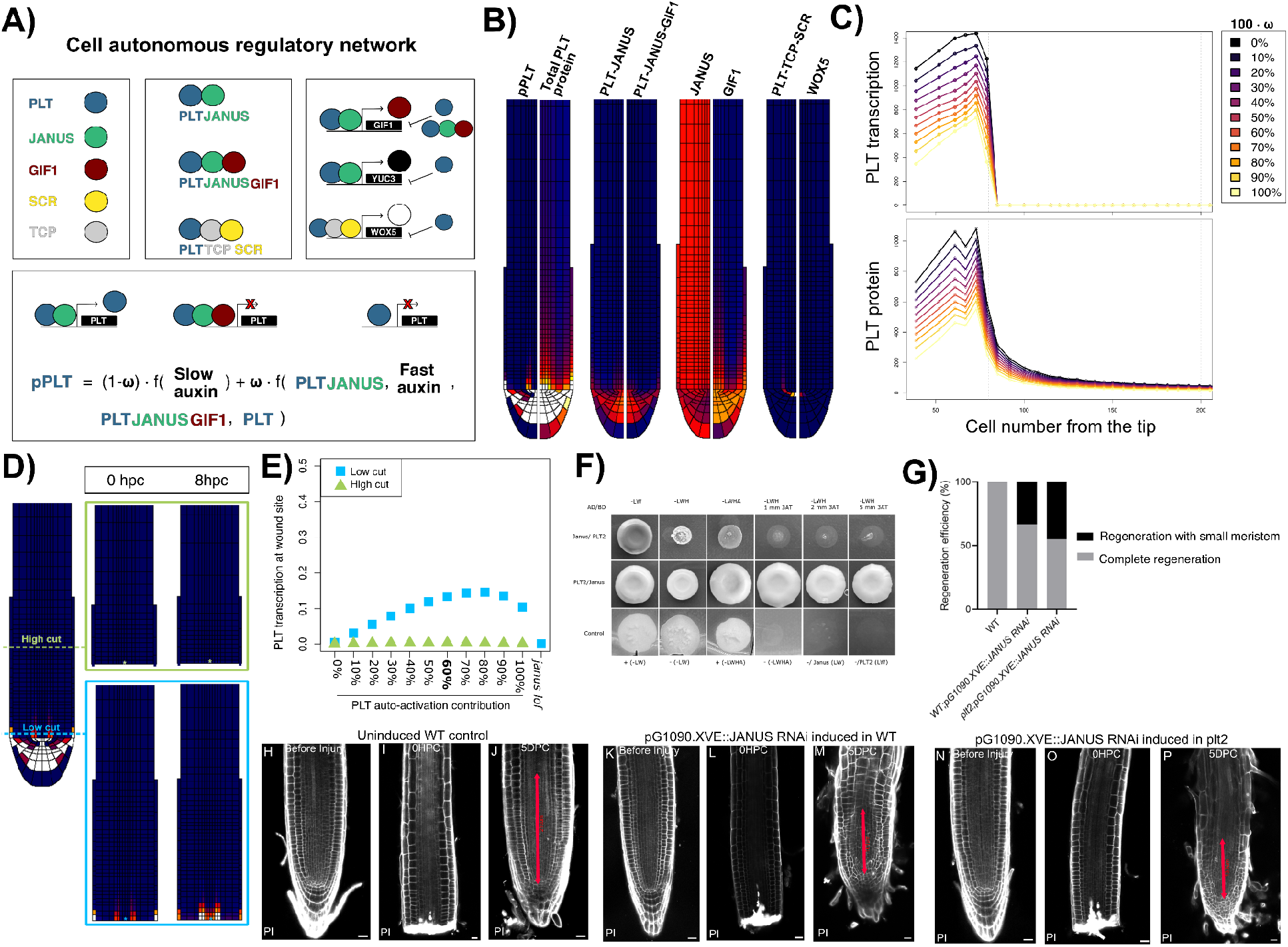
Auxin-PLT-JANUS-GIF1 model for PLT regulation in the root meristem. A) Cell autonomous regulatory network. PLT forms protein complexes to regulate the transcription of different targets, and unbound PLT represses them. PLT transcription is regulated slowly by auxin, and by PLT-JANUS and a fast auxin input. The contribution of each is defined by the parameter ω. PLT-JANUS-GIF and unbound PLT repress the transcription of PLT. B) Root model showing the emergent patterns of PLT, gene targets and protein complexes produced by the auxin-PLT-JANUS-GIF1 network. C) PLT transcription (top) and protein (bottom) levels along a longitudinal file of vascular cells at the centre of the root after 30 hours of simulation for different ω values. D) Root model showing PLT transcription prior, right after, and 8 hours after a LC or HC. Blue and green indicate the position where each cut is performed. (E) Quantification of PLT transcription in the vascular cell closest to the wound 8 hours post cut, for HC and LC for different ω values (PLT auto-activation contribution), and for the *janus* loss of function mutant. pPLT is normalized by the maximum production level enabled by the model parameters. (F) Reciprocal Y2H assays were performed using PLT2 and JANUS fused to the GAL4 DNA-binding (BD) or activation (AD) domains. Growth on selective medium lacking indicated Aminoacids, including medium supplemented with 3-AT. Controls included: positive (+) and negative (-) controls on -LW and -LHWA, PLT2BD and JANUS BD showed no autoactivation on -LW. G) Regeneration efficiency for WT, and an inducible RNAi line targeting JANUS in the WT and the *plt2* background. (H) Root apex before, and after a cut for WT, and an inducible RNAi line targeting JANUS in the WT and the *plt2* background.

## Results

### Integrating PLT auto-activation in a realistic model of the root apex

To explore the role of PLT2 auto-regulatory loop in intact and regenerating root tips (Durgaprasad et al., 2019), we used as a starting point our previously developed model incorporating an anatomically realistic root tip topology, detailed auxin transport and root growth dynamics (Methods; Mähönen et al., 2014; Salvi et al., 2020; Rutten and Ten Tusscher *2021;* Figure 1A-B). We included a cell autonomous regulatory network model of the auxin-PLT positive feedback loop previously established (Santuari et al., 2016; Salvi et al., 2020; Rutten et al., 2021), with PLT expression also regulated by itself. In searching for the mechanism underlying PLT auto-activation, we found that PLT1 forms a complex with JANUS. PLT1 binds its own promoter, and JANUS recruits RNA Polymerase II to activate gene transcription (Xiong et al., 2020). GIF1 is a transcriptional co-regulator that binds JANUS and blocks the binding site of Pol II, thus effectively repressing PLT1-JANUS-mediated *PLT1* expression (Ercoli et al., 2018; Xiong et al., 2020). The expression of *GIF1* itself is positively regulated by PLT1 (Ercoli et al., 2018). Based on the protein sequence similarity of PLT1 and PLT2 (Galinha et al., 2007; Kerstens et al., 2025), we hypothesize that PLT2 auto-activation in regeneration is also controlled through interactions with JANUS and GIF1, creating a negative feedback regulation that maintains PLT auto-activation within bounds.

Yeast-1-hybrid assays and pDAP–qPCR experiments have shown that PLT2 directly binds an evolutionary conserved box in its promoter (Kerstens et al., 2025), leading us to postulate that the PLT2 transcription factor recruits the mediator JANUS and the transcriptional regulator GIF1 to its promoter (Xiong et al., 2019; Xiong et al., 2020). Hence, in our model we assume that unbound PLT, and the protein complexes PLT-JANUS and PLT-JANUS-GIF1 can all bind the *PLT* promoter. To account for promoter binding site competition, we defined that while PLT-JANUS promotes PLT production, both PLT-JANUS-GIF1 and unbound PLT repress PLT production (Figure 1B). As a SCN marker we included in our model WUSCHEL-RELATED HOMEOBOX-5 (WOX5) whose expression is regulated by the protein complex formed between PLT, Teocinte-branched cycloidea PCNA 20/21 (TCP hereafter), and SCARECROW (SCR) (Shimotohno et al., 2018). Thus, our updated model includes multiple layers of PLT regulation in the root apex, to account for the fact that PLT can be unbound, or form a complex with JANUS, JANUS and GIF1, or TCP-SCR.

To explore the relative contribution of the auxin and the PLT-JANUS regulation on PLT transcription, we varied the fraction of PLT production attributable to each regulatory input, represented by the parameter ω, while keeping the total PLT production rate constant (Figure 1A). The model recovered the SCN-focused expression of PLT, as well as the graded PLT protein distribution for all ω values tested (Fig. 1C, Supplementary Figure 1AB), in agreement with experimental data (Mähönen et al., 2014). This model behaviour was robust to variations in the relative contribution of PLT-JANUS versus auxin on PLT transcriptional regulation (Fig. 1C, Supplementary Figure 1A-B), and in the repressive strength for PLT-JANUS-GIF1 and PLT in free form (Supplementary Figure 1D). Furthermore, our computational model predicted how the different protein complexes modelled are distributed in the root apex (Figure 1B). PLT-JANUS is predicted to be enriched in the SCN, and PLT-JANUS-GIF1 follows a near-identical pattern, with only slightly lower levels (Figure 1B). The expression of GIF1 mirrors the PLT-JANUS complex with highest levels in the SCN area (Ercoli et al., 2018), a reflection of this complex promoting its expression. Conversely, unbound JANUS levels are low within the SCN and higher in the cells surrounding it. As experimentally described PLT-TCP-SCR is enriched in the cell layer surrounding the stele close to the SCN with a maximum in the QC, where WOX5 is expressed (Shimotohno et al., 2018).

Although loss of JANUS function causes embryo lethality (Xiong et al., 2019), our computational model enables us to explore how PLT regulation in the root apex would be affected by eliminating its activity (Methods). We found that this perturbation reduced PLT transcription (Supplementary Figure 1F-G), consistent with JANUS RNAi data (Xiong et al., 2020). Conversely, simulating *gif1* increased PLT transcription in the SCN, consistent with PLT1 observations (Ercoli et al., 2018), indicating that GIF1 maintains in check the PLT autoactivation regulation mediated by PLT-JANUS (Xiong et al., 2020).

To test the key assumption made in our model related to PLT-autoactivation, we investigated the binding between PLT2 and JANUS proteins. We performed a yeast two hybrid (Y2H) assay and confirmed the direct interaction between PLT2 and JANUS in both reciprocal BD/AD configurations (Figure 1G). Altogether, our auxin–PLT–JANUS–GIF1 model integrates the roles of auxin and PLT–JANUS regulation in controlling PLT transcription, and recapitulates the PLT expression in WT, *janus*, and *gif1* Arabidopsis roots.

### PLT–JANUS Complex Formation Drives Primary Root Regeneration

Next, we used our auxin-PLT-JANUS-GIF1 model to simulate root regeneration experiments. We defined the cut position as either just above the QC (low cut-LC, blue), or at the distal end of the meristem (high cut-HC, green), and removed all cells below the cut from our simulation domain (Figure 1D; Methods). To infer regeneration competence, we focus on the build-up of PLT transcriptional levels in a vascular cell next to the cut site after 8 hours post-cut (hpc) (Durgaprasad et al., 2019). HC simulations failed to produce a PLT transcriptional response regardless of the contribution of PLT autoactivation (ω) (Figure 1DE, and Supplementary Figure 1C). By contrast, all LC simulations with non-zero PLT autoactivation produced a response at 8hrs post-cutting. Thus, our auxin-PLT-JANUS-GIF1 model reproduces the persistent PLT protein levels associated with successful regeneration in LC, and the regeneration failure observed in HC (Durgaprasad et al., 2019).

Because both auxin signalling and PLT-JANUS regulate PLT expression, we used our model to assess their individual role in the regeneration response. We simulated a LC in which PLT production was regulated exclusively by auxin, and found that this was insufficient to elicit a rapid PLT transcriptional response in vascular cells near the wound (Supplementary Figure 1E). Conversely, incorporating only the PLT-JANUS-mediated regulation produced a rapid response on its own, although less pronounced than when both auxin and PLT-JANUS are involved (Supplementary Figure 1E). This result indicates that auxin alone is insufficient and that PLT autoactivation is required for the early regeneration response, supporting that they function as a single integrated regulatory module controlling regeneration. Next, we simulated a LC in our model of the *janus* mutant and found that in this case the root stump produced no PLT transcriptional response within 8 hpc (Figure 1E), suggesting a defect in regeneration. Moreover, this result indicated that the residual PLT proteins in the in-silico root stump kickstart PLT transcription through the formation of the PLT-JANUS complex, and otherwise it will fail.

Next, we seek to validate the crucial role of PLT and JANUS on regeneration. Previously it was shown that the plt2 mutant does not show any defect in root regeneration (Durgaprasad et al., 2019). To explore the role of JANUS we performed regeneration frequency assays using an estradiol-inducible line *pG1090i::JANUS-RNAi* (Methods). This inducible line was used because the loss-of-function of JANUS is embryo lethal (Xiong et al., 2019), and continuous induction of RNAi occurred from 3 days before the cut until the end of the experiment. We performed a cut at the position of the cells above the QC (LC) and found that while all the resected root apices regenerated, 33.33% of induced WT root tips failed to restore the original meristem length (Figure 1 K-M, Q), unlike the uninduced controls (Fig 1 H-J, Q). As modelling data shows that PLT2-JANUS interaction is necessary for successful regeneration, we asked whether a reduction in JANUS transcripts in the *plt2* mutant background will produce any effect on root tip regeneration. Strikingly *plt2*; *pG1090*.*XVE::JANUS RNAi* root tips regenerated, but 45% of the root tip failed to restore the original meristem length (Fig 1 N-Q). The even higher frequency of plants with a partially restored meristem in the absence of both JANUS and PLT2, underlines their joint role in root regeneration. This result further reinforces our finding that PLT2 interaction with JANUS is essential for regeneration.

### PLT Overexpression Disrupts the PLT Regulatory Balance

Next we used our model to simulate the exogenous induction of PLT produced by an overexpression and DEX-inducible nuclear localization system, *35S::PLT2-GR* (Durgaprasad et al., 2019). To do so, we included in our model a DEX-driven factor for inducible and constitutive PLT production (Methods) and simulated 50 hours of DEX treatment. Below, we compare the response in a cell just above the QC cells, and another cell in the upper half of the meristem (Figure 2A).

**Figure 2.**
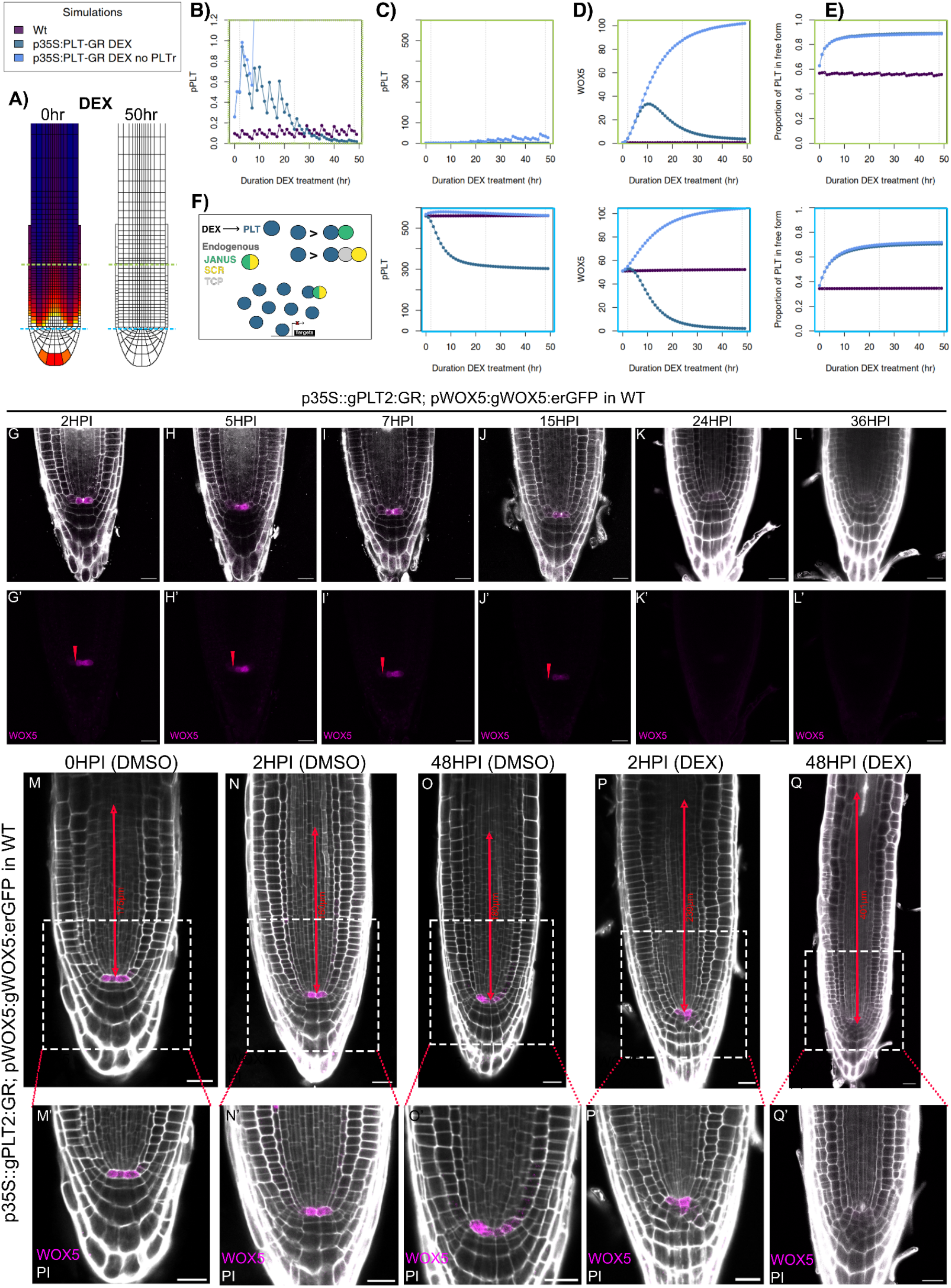
Non-monotonical effects of DEX-induced PLT accumulation. (A) PLT protein before and after 50 hours of DEX-induction of PLT. Temporal effects of 50 hours of DEX induction on cells at two positions of the in-silico root apex (see blue and green coloured lines in (A)): (C) PLT transcription (closeup shown in (B)), (D) WOX5, and (E) the proportion of PLT in free form. Notice how for *p35S::PLT-GR* the PLT-targets are first up-regulated and then down-regulated, and how the response is not observed in the *p35S::PLT-GR* without the repressive role of PLT in free form (‘no PLTr’). Dashed grey lines indicate 0, 2h or 48h of DEX treatment. (F) Diagram illustrating how the very high levels of PLT in free form competes with PLT-JANUS resulting in the reduction in expression of PLT targets. (G-L) Expression of a WOX5 reporter after different times of DEX induction in the p35S::PLT2-GR line. The red arrow in the lower panels marks the expression of WOX5 in the cells of the root apex. (M-Q) *WOX5* expression in the *p35S:PLT2-GR* inducible line after at 0, 2, and 48 hours of treatment with either DEX or DMSO.

As expected, we found that the total PLT protein levels (unbound and in complex) gradually increase with the duration of the DEX treatment (Figure 2A; Supplementary Figure S2). The trend with which PLT levels increase is independent of the position along the root tip. In contrast, the response of the endogenous PLT targets markedly differed along the root apex and were highly non-monotonic (Figure 2B-D). Above the QC, PLT transcriptional levels (pPLT) dropped to approximately half of WT levels (Figure 2C). The remaining expression corresponds to the auxin-driven part of PLT transcription. In the cells of the proximal meristem, where *PLT* is normally not expressed, DEX treatment led to a transient and minor induction of *PLT* transcription, before returning to zero (Figure 2B and C). In the case of WOX5, the QC cells show a reduction in its expression to negligible levels after 30 hours of DEX treatment (Figure 2D). In the proximal endodermal meristematic cells *WOX5* showed a mild upregulation followed by a drop at longer DEX induction times (Figure 2D). Thus, our simulations predicted a temporal non-monotonic regulation of *PLT* and *WOX5* expression in cells at different locations within the root meristem. Importantly, these non-monotonic responses were not recovered in a version of our model where PLT auto-activation is mediated directly through free PLT (Supplementary figure S3), thus highlighting the role of the PLT-JANUS-GIF1 complexes in this response.

We set to experimentally test the model’s prediction that prolonged DEX treatments will cause a mild expansion, and then the complete disappearance of *WOX5* in the root apex. To this end we crossed the DEX-inducible steroid inducible transgenic line *35S::PLT2-GR*, with a translational *WOX5* reporter *pWOX5::gWOX5:erGFP*. We monitor the WOX5 reporter throughout 48 hours of DEX induction (Figure 2G-Q). We noticed that the activity of the WOX5 reporter gradually decreased from 15 hours of DEX treatment onwards, until it was undetectable (Figure 2G-L). Importantly, the lack of *WOX5* expression after 48 hours was only produced in DEX but not DMSO treated plants (Figure 2M-Q), confirming the prediction of our computational model (Figure 2D).

Next we aimed to use our model to understand the mechanistic basis of the drop in expression WOX5, and of PLT. The model predicted that, after 48 hours of DEX induction, PLT–TCP–SCR and PLT–JANUS levels would be high throughout the meristem (Supplementary Figure S2). These two protein complexes could in principle induce the expression of their target genes, yet our model predicts their transcription actually decreases after prolonged induction (Figure 2B and C; Supplementary Figure S2). This finding suggests that further increasing PLT levels compromises the capacity of the PLT protein complexes to regulate their target genes. We analysed the proportion of unbound PLT relative to the total PLT levels in our model, (Figure 2E) and noticed that the longer the DEX treatment, the more PLT was in free form. As unbound PLT can bind the PLT promoter on its own (Kerstens et al., 2025), we reasoned that such increase causes the downregulation of PLT and WOX5 due to competition with PLT-JANUS and PLT-SCR-TCP, respectively. In agreement with our hypothesis, simulations of DEX induction without the repressive effect of unbound PLT (‘no PLTr’) produced a continuous positive response on *PLT* and *WOX5* throughout 50 hours of DEX induction (Figure 2 and Supplementary Figure S2).

Thus, our simulations reveal a non-monotonic response of PLT targets in DEX induction experiments due to the very high levels of PLT protein shifting the balance between complex-bound protein and unbound protein, resulting in the repression of PLT targets by PLT (Figure 2F).

### Non-monotonical Regeneration Competence of High Cuts

Our DEX mediated PLT-induction simulations predict that the competence of PLT to regulate its target genes is non-monotonic, first inducing their ectopic expression and later repressing it (Figure 2). We hypothesize that this response accounts for the paradoxical, non-monotonic improvement in HC regeneration competence following different durations of DEX induction in *p35S::PLT2-GR* (Durgaprasad et al., 2019).

To test this hypothesis, we simulated a LC and a HC after different durations of DEX treatment prior to the cut (Figure 3). We ran the simulations for 8 hpc and assessed regeneration competence by comparing the PLT transcriptional response in each simulation with that of control LC simulations without DEX induction. We defined the PLT transcription level reached in these control simulations as the minimum level required for regeneration competence (Supplementary figure S4B). The model recovered that a short DEX induction treatment was sufficient to confer regeneration competence to HC simulations; however, competence declined with longer treatment durations (Figure 3A-B; Supplementary Figure S4). LC simulations were regeneration-competent even before a DEX treatment, but exhibited a similar decline in competence with increasing DEX-exposure duration (Figure 3A-B; Supplementary Figure S4). These results are consistent with the regeneration competence of HC experimentally observed in roots (Figure 3C; Durgaprasad et al., 2019). Removing the repressive role of unbound PLT on PLT expression in our simulations, produced a continuous positive effect of PLT induction on regeneration competence (Supplementary Figure 4C). Therefore, similar to our simulations in intact roots (Figure 2), this counterintuitive non-monotonic regeneration competence response is caused by an early increase in PLT levels enabling sufficient PLT-JANUS complex to trigger auto-activation of PLT, and further increments reducing the ratio of PLT-JANUS to unbound PLT.

**Figure 3.**
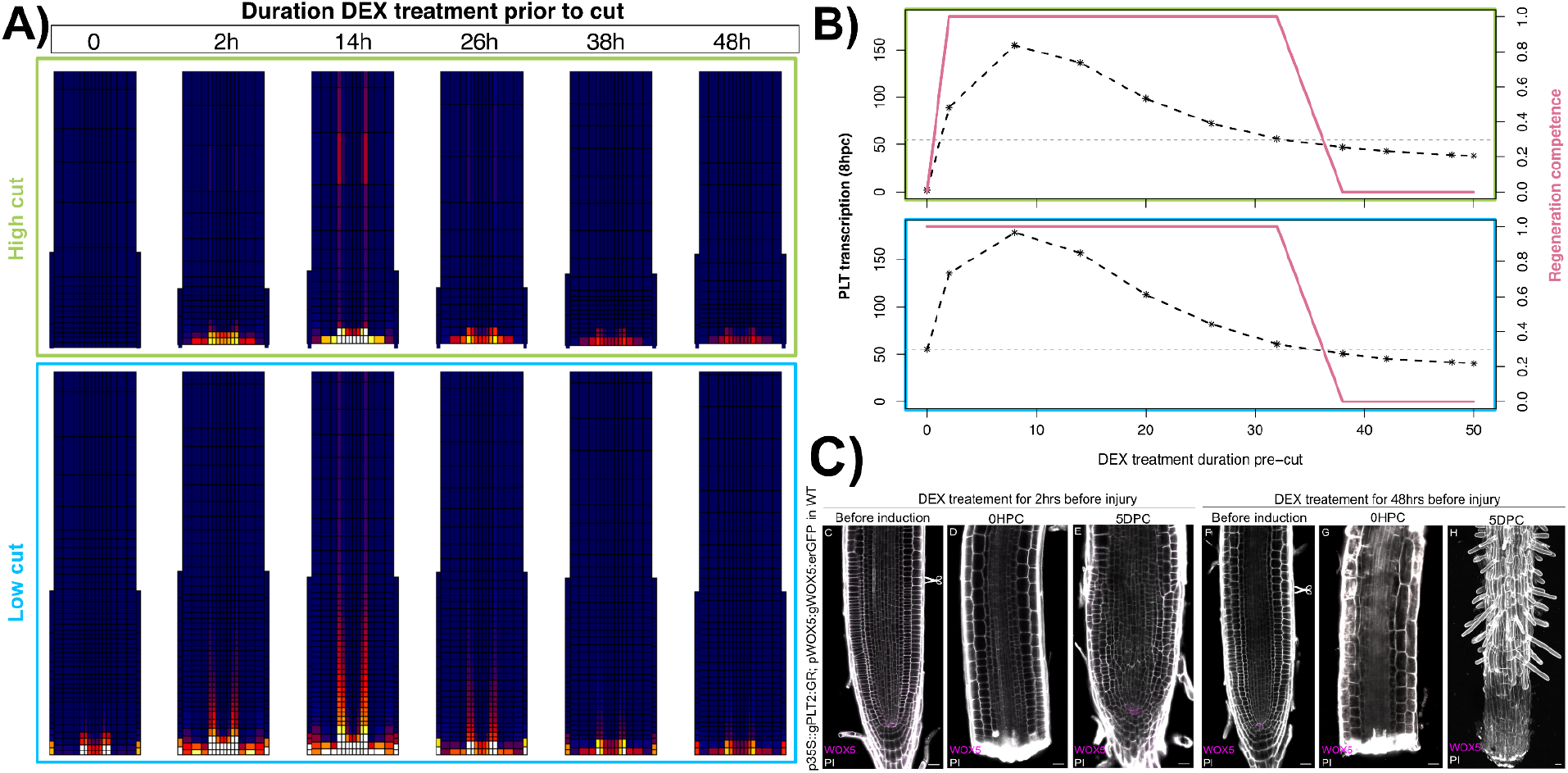
DEX pre-treatments produce a non-monotonic response in PLT expression and regeneration competence. (A) PLT transcription 8 hours post-cut in root apices treated with DEX prior to high (green) or low (blue) cuts. (B) Quantification of PLT transcription in the vascular cell closest to the wound after 8 hours after cut, in HC and LC. Pink indicates regeneration competence of the in-silico root stumps based on whether PLT transcription (black dashed line) is equal or higher than a threshold (horizontal grey line). (C) Experiments showing regeneration or differentiation of *p35S::PLT2-GR* root tips upon removal of cells outside the competence zone. Short-term DEX treatments (2 hours) result in root tip regeneration, whereas long-term induction (48 hours) result in differentiation and failure to regenerate.

Thus, our auxin-PLT-JANUS-GIF1 model recovers the temporal window in which the PLT induction produces PLT levels where autoactivation, and hence regeneration competence, are effective, and predicts that accumulation of unbound PLT terminates the response.

### Co-induction of PLT and an interacting protein extends regeneration competence in-silico

Based on our findings, we hypothesize that co-induction of both PLT and a PLT interacting protein could counteract the negative effects produced by prolonged induction of PLT nuclear localization, by maintaining an adequate ratio of unbound PLT to complex-bound PLT. We used our model to explore the plausibility of this hypothesis and simulated the induction of either JANUS or SCR together with PLT (Figure 4A).

**Figure 4.**
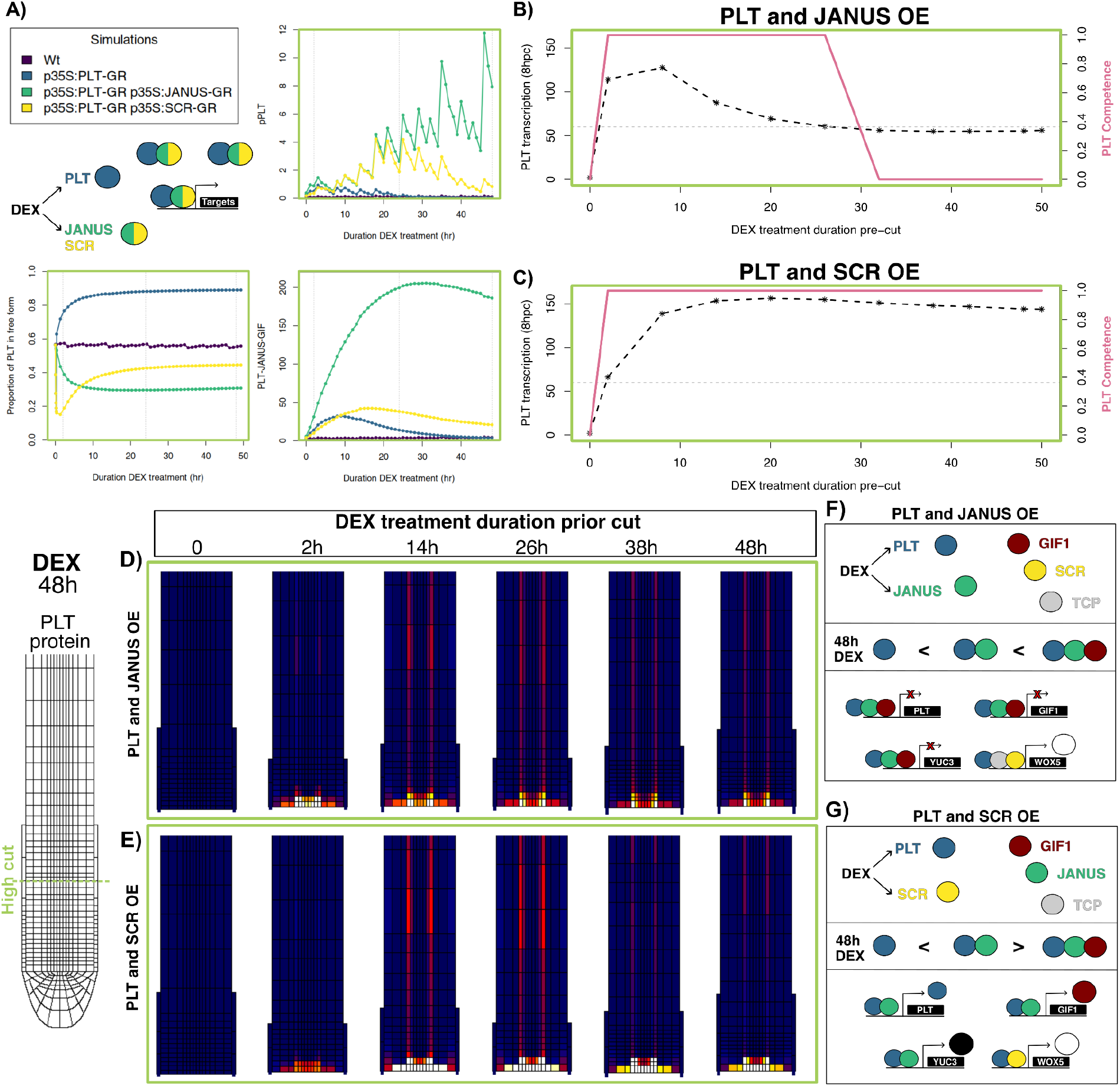
PLT2 and JANUS or SCR co-induction simulations. (A) The simulation of DEX-induction of both PLT and an interacting partner (JANUS or SCR) and their effects on PLT expression, the proportion of PLT in free form, and total PLT-JANUS-GIF1 complex in the cells at the end of the meristem. Notice that both JANUS and SCR co-induction with PLT result in low PLT free levels compared to induction of PLT alone, thus representing two alternative ways of maintaining free PLT levels in check. HC were performed after different durations of DEX treatment for co-induction of PLT and JANUS (B and D), or of PLT and SCR (C and E). (B) and (C) show PLT transcription in a vascular cell close to the wound 8 hours post-cut in roots treated with DEX for different durations, and their competence (threshold shown as a horizontal grey line). (D-E) PLT transcription in the root model 8 hours after a high cut. Schemes illustrating the effects on PLT targets after 48 hours of PLT induction with either JANUS (F) or SCR (G).

As expected, co-induction with JANUS produced much higher levels of PLT-JANUS complex, while SCR co-induction produced higher levels of PLT-TCP-SCR (Supplementary Figure S5). Notably, in both cases the ratio of PLT in free form decreased relative to the induction of PLT alone, thus allowing for the regulation of PLT targets (Figure 4A). Interestingly, the co-induction of PLT and SCR still produced subtle detrimental effects on the expression of PLT-complex targets as can be seen for *WOX5*, whose expression in a cell at the end of the meristem follows a decreasing trajectory (Supplementary Figure S5). This can be understood from a combination of TCP levels being limiting and PLT free levels still being high relative to the non-induced condition (Supplementary Figure S5).

Next, we performed HC simulations after different durations of DEX co-induction of PLT with either JANUS or SCR (Supplementary Figure 6AB). Surprisingly, we found that the long-term co-induction of PLT and JANUS still resulted in a drop in PLT transcription at the wound site at 8hpc, just below our regeneration competence threshold (Figure 4B). This result suggests the regeneration success of PLT and JANUS co-induction after prolonged DEX induction will be mildly increased compared to the induction of PLT alone (Figure 3B). Contrarily, PLT and SCR co-induction maintained high PLT transcriptional levels and regeneration competence, regardless of the duration of DEX treatment prior to the cut (Figure 4C). The stark difference between JANUS and SCR co-induction simulations relate to GIF1 being one of PLT-JANUS targets, as this sets a break in PLT-JANUS regulation (Figure 4F). Indeed, PLT-JANUS-GIF reaches extremely high levels in the PLT and JANUS co-induction simulation (Figure 4A). In agreement with this interpretation, a PLT and JANUS co-induction simulation without PLT-JANUS-GIF1 repressing PLT transcription resulted in sustained regeneration competence (Supplementary Figure S6CD). Importantly, our simulations indicate that these outcomes are robust to variation in expression of the second protein (Supplementary figure 6E).

Thus, our simulations predict that co-induction of PLT with a PLT-interacting protein can enhance regeneration competence, with the identity of the co-induced partner determining the outcome: sustained regeneration response with SCR, or a modest improvement with JANUS.

### SCR and JANUS overexpression titrates excess free PLT to maintain root regeneration competence *in planta*

Next, we sought to validate our simulation results by testing the hypothesis that co-induction of PLT with a PLT-binding protein can further extend regeneration competence following a HC. We generated transgenic lines for co-induction of PLT2 with either JANUS (WT/p35S::PLT2-GR; p35S::JANUS-GR) or SCR (WT/p35S::PLT2-GR; pRPS5A::SCR-GR) (Figure 5A). We assessed transcript levels in these co-induction lines to confirm that both transgenes were expressed at comparable levels (Figure S7). We next examined root tip regeneration in these co-induction lines. Consistent with previous reports, induction of PLT2 alone for 48 hours did not promote root tip regeneration following a HC (Durgaprasad et al., 2019; Figure 5B-E). In contrast, co-induction of PLT2 with either JANUS or SCR for 48 hours resulted in faithful restoration of the root tip following a HC (Figure 5B, F-K), although the regeneration efficiencies differed between the two co-induction lines (Figure 5B). In line with the predictions from our model, the co-induction of PLT with SCR produced a higher regeneration response compared to JANUS (Figure 4BC and 5B).

**Figure 5.**
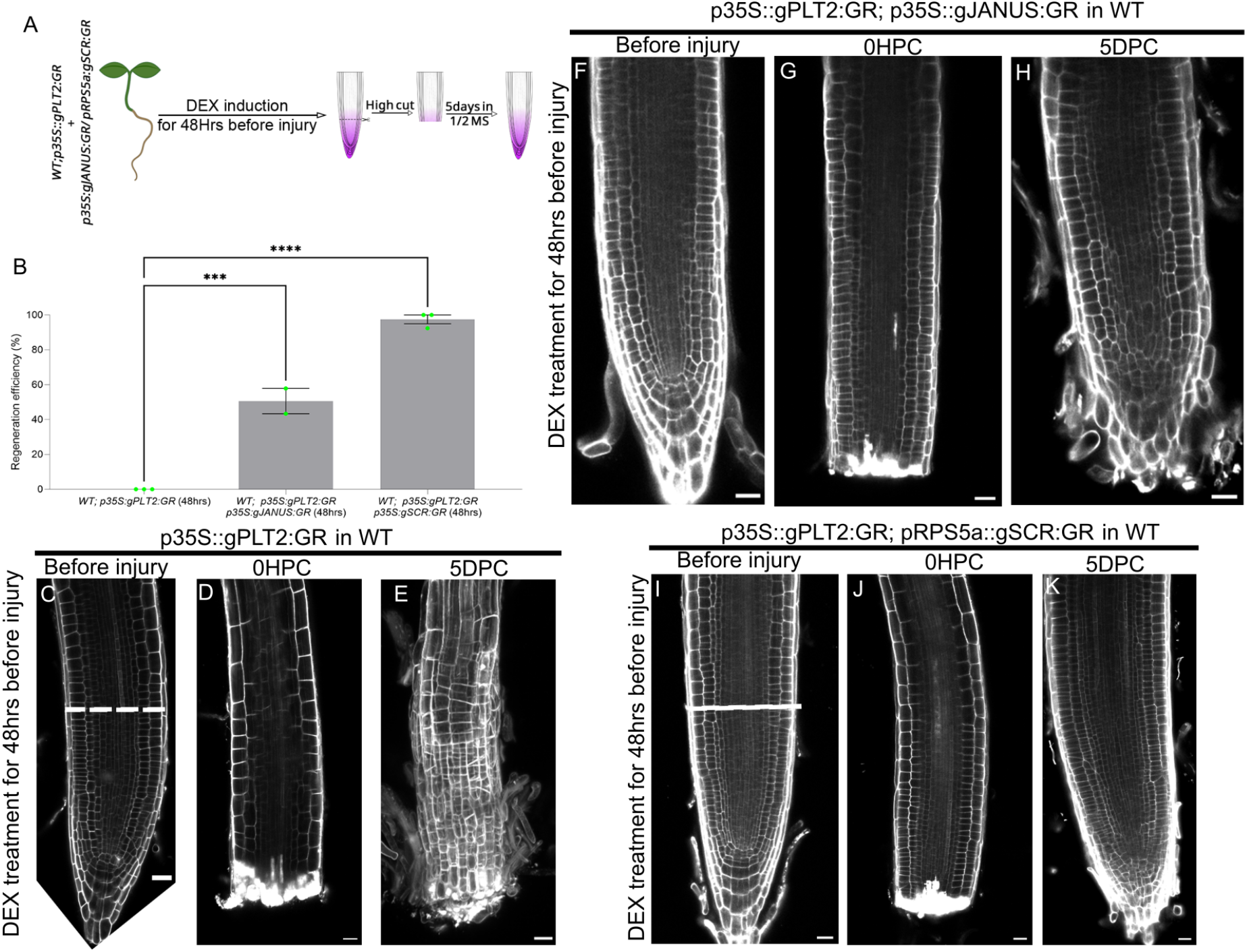
Co-induction of PLT and an interacting protein can extend regeneration competence in HC. (A) Experimental setup showing DEX induction for 48 hours, and then a HC is performed. (B) Regeneration efficiency of HCs when inducing PLT2, PLT2 and JANUS, or PLT2 and SCR for 48 hrs of DEX treatment prior to the cut. Time lapse confocal images showing root apex after 48 hours of DEX induction prior to the cut, right after the cut, and 5 days post cut for *p35S::PLT2-GR* (C-E), *p35S::PLT2-GR p35S::JANUS-GR* (F-H), and *pRPS5a::SCR-GR* (I-K).

Together, these experimental results validate our model predictions and support our proposal that co-induction of a PLT-binding protein maintains a larger pool of PLT in the complex-bound state, thereby sustaining regulation of PLT target genes and promoting successful root tip regeneration.

## Discussion

The dosage of the PLT transcription factors is a key determinant of developmental decisions in plants, including the establishment of root zonation (Mähönen et al., 2014), the development of lateral root primordia (Du et al., 2017), embryogenesis (Kerstens et al., 2025), and regeneration from tissue explants (Kareem et al., 2015) and severed organs (Durgaprasad et al., 2019). In intact roots the localized PLT expression in the root stem cell niche is regulated by a slow auxin-mediated transcriptional induction (Mähönen et al., 2014), whereas regeneration after root tip resection is driven by a faster PLT auto-activation transcriptional response (Durgaprasad et al., 2019). Previous work indicated that for this autoactivation loop to successfully induce regeneration, the extent of root excision and the resulting lingering PLT levels are a key factor (Durgaprasad et al., 2019). Still, several major questions remain.

Firstly, how are slow auxin mediated induction and faster PLT mediated auto-activation integrated during normal root development? Secondly, how come a short PLT induction enhances whereas sustained PLT induction reduces regeneration potential (Durgaprasad et al., 2019). Together, these observations suggest the involvement of additional regulatory layers in how PLT levels are translated into transcriptional and developmental responses.

Here we put forward a computational model of a root apex that incorporates the role of slow auxin-mediated and a fast PLT auto-activation in PLT expression, showing they form an integrated regulatory module operating in intact roots, and in root tip regeneration (Figure 1). Our model explicitly accounts for the formation of different PLT protein complexes, and the distinct regulatory activities of free and complex-bound PLT in regulating the expression of its target genes (Figure 1B). Through our computational-experimental approach we revealed a ratiometric balance between bound and unbound PLT proteins for the regulation of PLT targets. Specifically, PLT and WOX5 expression require the PLT-JANUS and PLT-TCP-SCR complexes relative to unbound PLT, which would otherwise inhibit their expression (Figure 2). Furthermore, we show that this ratiometric balance underlies the control of regeneration. Roots in which most meristematic cells are excised (HC) will regenerate following a short induction of PLT accumulation prior to the cut, owing to the accumulation of PLT-JANUS relative to free PLT allowing for PLT expression. In contrast, further induction of PLT shifts the ratiometric balance towards unbound PLT, leading to a decrease in *PLT* and *WOX5* expression (Figure 2), and preventing regeneration of the root apex (Figure 3). Thus, our results demonstrate that rather than responding solely to absolute PLT concentration, PLT targets respond to the relative enrichment of complex-bound versus unbound PLT.

Based on the uncovered relationship between PLT dosage, the protein complex balance and regeneration competence, we hypothesized that prolonged induction of PLT together with an interacting protein could further extend the regeneration competence of HCs. In particular, our computational model predicted that co-inducing PLT with SCR will produce a sustained positive effect on regeneration competence (Figure 4C) compared to co-induction with JANUS (Figure 4B). This difference arises from the GIF1-mediated negative feedback, which interferes with PLT-JANUS and represses PLT expression independently of the ratiometric balance. Our experimental validation of these predictions showed a higher regeneration frequency of the PLT and JANUS co-induction line compared to the control (PLT induction alone), and the highest regeneration observed in the PLT and SCR co-induction line (Figure 5). Thus, these results support the conclusion that maintaining the ratiometric balance between PLT–JANUS complexes and unbound PLT are necessary for PLT–JANUS-mediated gene expression and regeneration.

Previous work showed that MIR396 overexpression promotes regeneration of HC (Baulies et al., 2025), consistent with miR396 acting as an indirect positive regulator of PLT (Rodriguez et al., 2015). Nevertheless, the regeneration competence induced by constitutive overexpression of MIR396 differs from the non-monotonic response observed in our PLT induction experiments (Baulies et al., 2025; Durgaprasad et al., 2019; Figure 3). Importantly, these results are only seemingly contradictory. In the case of MIR396 induction, the downstream increase in PLT remains subject to the negative feedback through GIF1 regulatory loop. Consequently, the increase in PLT remains within bounds resulting in a high PLT-JANUS to unbound PLT ratio, and sustained regeneration competence. In contrast, in our experiments the exogenous PLT accumulation is produced through an inducible PLT2 overexpression system (*p35S::PLT2-GR*) and is therefore independent of GIF1 regulation. This system allows us to explore the consequences of PLT accumulation beyond its physiological range.

The lower regeneration benefit of the co-induction of PLT with JANUS compared to SCR shows that not just every interacting protein can maintain the ratiometric balance (Figure 4), owing to the connectivity of each protein complex in the PLT regulatory network. These results underscore the importance of computational models for unravelling how different protein complexes are integrated within the regulatory networks that control target gene expression (Strotmann et al., 2025). Our results thus indicate that the various PLT-protein complexes not only allow for diversification of the regulatory targets of this master regulator of root development, but also enable sophisticated tuning of positive and negative feedback effects, balancing the initial boosting and subsequent curtailing of positive feedback.

Together, our computational modelling–driven approach, coupled with experimental validation, demonstrates that PLT dosage is not a linear readout. Instead, the regulatory network incorporating protein-complex formation converts quantitative changes in PLT levels into non-monotonic transcriptional responses. The ratiometric mechanism uncovered here is therefore a key element of the complex regulatory landscape through which PLT proteins establish rapid responses during regeneration. This mechanism may also play a role in initiating germination (Salvi et al., 2020) as well as the stable maintenance of stem cell niches (Mähönen et al., 2014; Du et al., 2017).

## Methods

### Yeast two-hybrid (Y2H) Whole CDS

Protein–protein interaction assays were performed using the GAL4 System as described (De Folter & Immink, 2011). The full-length coding sequence of JANUS was amplified from cDNA and cloned into pDONR221 entry vectors and subsequently into the pDEST22 and pDEST32 expression vectors using the Gateway™ system. Expression vectors for PLT2 were previously generated (Kerstens et.al 2025). Expression vectors were transformed into the PJ69-4A (pDEST22) and PJ69-4α (pDEST32) yeast strains, and the different combinations were acquired by mating. Reciprocal Protein–protein interactions were screened on -LW, -LWA and -LWH dropout medium, supplemented with 1, and 2mM 3-amino-1,2,4-triazole (3-AT). Plates were incubated for 5 days at 20°C. Auto activation tests were performed using solely PLT2BD and JANUSBD on -LWHA. For positive and negative controls of previously tested proteins have been used.

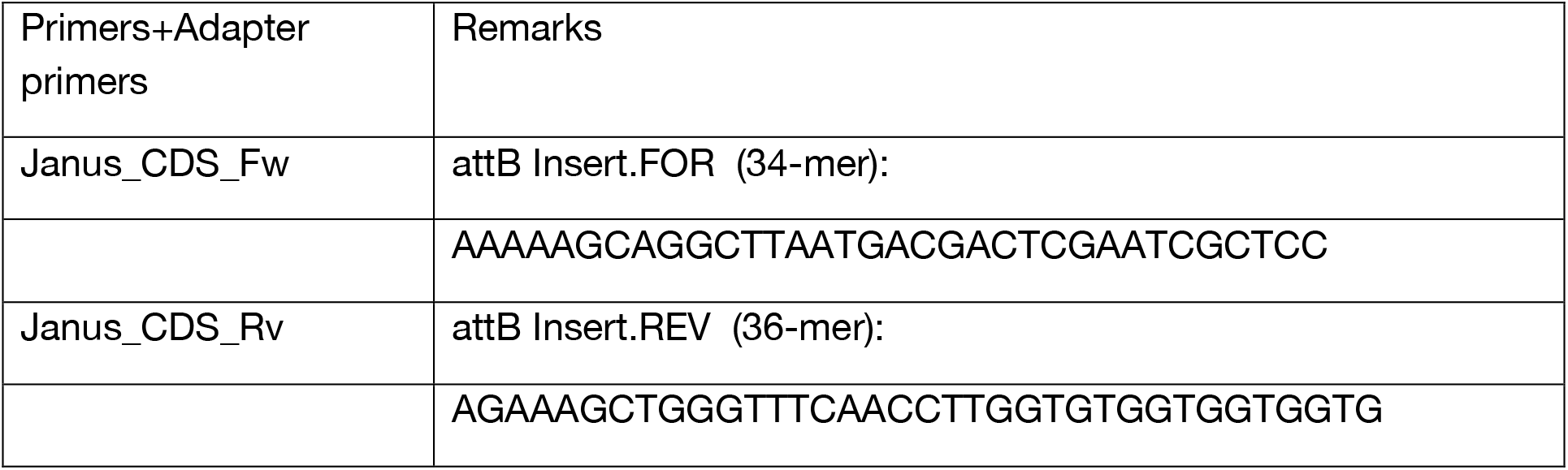

### Regeneration assay

Root tip resection and the regeneration assay were performed as previously described (*Durgaprasad et al., 2019*). The regeneration assay was done on 4-day-old seedlings. The seedlings were placed on a sterile glass slide with few drops of sterile water. The root tips were resected using a sharp 1ml syringe tip (Dispovan) under a Zeiss AxioScope A1 microscope. The resection was performed within the root meristem, and the resected section was measured to ensure high / low cuts. The high and low cuts were defined by the distance from the fourth tier of the columella cells to the cut site. Low cuts were between 120 and 210μm, whereas high cuts were between 250 and 300μm. Post resection, the seedlings were prepared for confocal imaging following returning to MS medium. The root tip regeneration was evaluated based on the growth and morphology of the root tip at 5-days post-cut (DPC).

### Treatments

#### Estradiol treatment

In case of pG1090i::JANUS RNAi in WT and *plt2*, 1dpg old seedlings were transferred to half strength MS medium supplemented with 5μM estradiol and were incubated till resection. Post resection, the cut roots were placed on fresh half strength MS medium supplemented with 5μM estradiol till 5DPC (to give continuous induction). In case of control, the 1dpg seedlings were transferred to half strength MS medium supplemented with DMSO, before resection.

### Dexamethasone treatment

In case of p35S::gPLT2:GR, p35S::gJANUS:GR in WT and p35S::gPLT2:GR, pRPS5a::gSCR:GR in WT 2dpg old seedlings were transferred to half strength MS medium supplemented with 20μM dexamethasone (DEX) and were incubated for 48hrs before resection. Post resection, the cut roots were placed in fresh half strength MS medium till 5DPC. For control MS medium supplemented with DMSO was used instead of DEX. In case of 2hr induction, 4dpg seedlings were transferred to half strength MS medium supplemented with 20μM DEX and incubated for 2hrs before resection. Following resection, the cut roots were placed in fresh half strength MS medium till 5DPC.

### Computational model

We developed a model of a root apex using as a basis earlier multi-scale root models using a realistic root tip architecture, polar auxin transport mediated by PINs, auxin influx regulated by AUX1 (Salvi et al., 2020; Rutten and Ten Tusscher, 2021). The model implements realistic cell growth and division dynamics in the root apex. Namely we define the position of the different root zones, with a stem cell niche overlaying the columella where no growth or division takes place, followed by a meristematic zone where cells actively grow and divide; upon reaching the border to the elongation zone where cells grow rapidly and anisotropically. The model also incorporates a cell type arrangement representing the concentric cylinders found in the root, with the narrow stele cells at the centre, followed by the pericycle, endodermis, cortex, epidermis and lateral root cap. The different cell layers have predefined properties regarding the expression and polarity of the auxin efflux transporters PINs (Mähönen et al., 2014), influx transporters AUX1 and LAX3, and plasmodesmata density patterns (Rutten and Ten Tusscher, 2021). An auxin influx from the top represents shoot-derived input, and auxin can also be produced by the cell autonomous regulatory network (see details below).

### Cell autonomous regulatory network

Each cell in the simulation has a regulatory network that models the auxin signalling pathway, the expression of PLT, JANUS, GIF1, WOX5, YUC3, PIN, AUX1 and LAX3, and the association and dissociation of different PLT protein complexes. The details of each subpart of the model network are explained below.

#### Auxin signalling regulation

The auxin signalling pathway is modelled as previously (Mähönen et al., 2014; Salvi et al., 2020). Namely, we model generic AUX/IAA and ARF factors, with the auxin levels of the cell modifying the fraction of ARF that is unbound of AUX/IAA pathway repressors; this free ARF fraction can regulate the gene expression of auxin targets. We assume that auxin regulation of AUX/IAA degradation is fast (Mähönen et al., 2014), and used a quasi-steady-state assumption (QSSA) to derive the AUX/IAA levels in each cell:

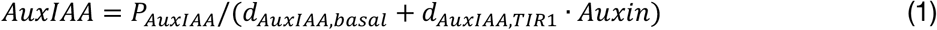

Each cell has a constant amount of ARF, and we use the AUX/IAA expression (Equation 1) to calculate the amount of ARF that is in free form:

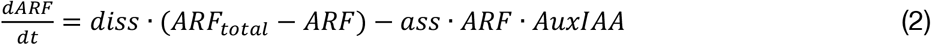

To account for the slow regulation of PLT transcription, important for separating the root zonation patterning from fast auxin responses, i.e. tropisms (Mähönen et al., 2014), we included A, B, C and D as four intermediaries mediating the regulation from ARF to PLT transcription. An unintended effect of this cascade is that each subsequent intermediary regulator has lower activity and thus does not reflect the auxin input. To correct for this effect, we used *α*_1_ as a correcting factor in the production of each intermediary as previously (Salvi et al., 2020). This term ensures that each intermediary reaches activity levels that align with the original high auxin input:

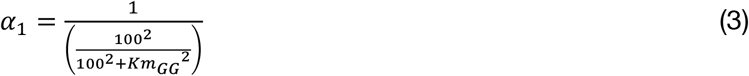

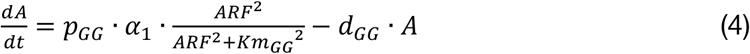

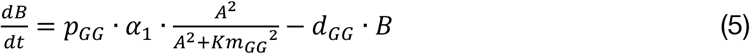

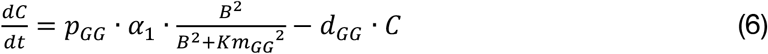

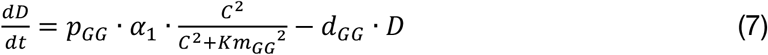

These intermediaries allow us to define *AuxinR*1 and *AuxinR*2 to describe slow and fast auxin signalling regulation of PLT regulation, respectively:

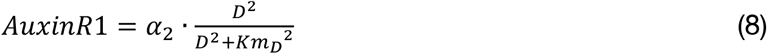

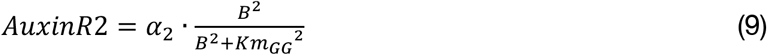

As for the auxin intermediaries, we used a correcting factor (*α*_*2*_) to ensure that PLT regulation aligns with the original auxin input:

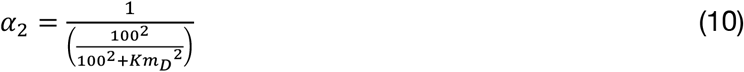

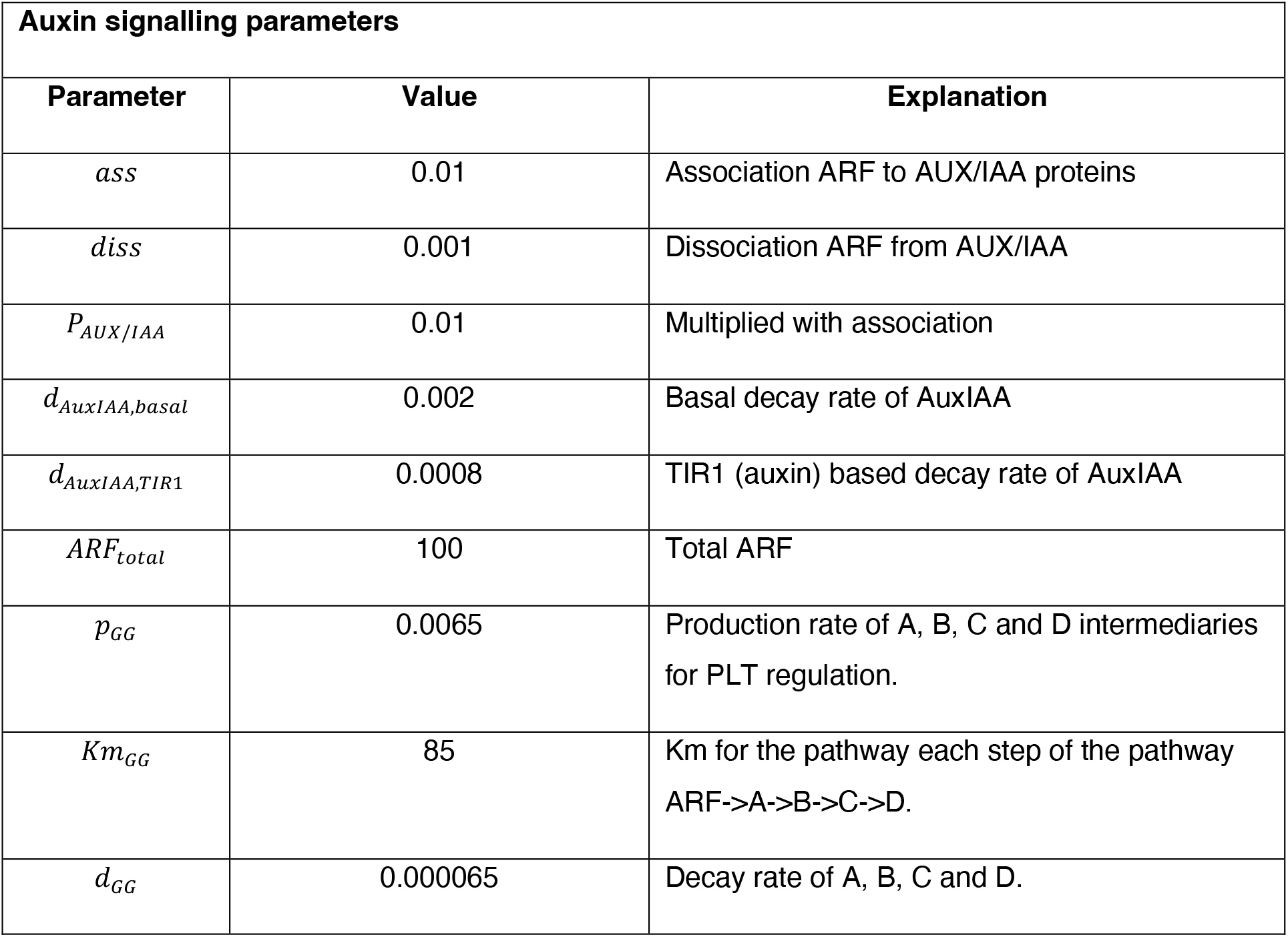

### PLT regulation

The effect of slow auxin signalling on the regulation of PLT transcription has been studied and modelled before (Mähönen et al., 2014; Salvi et al., 2020; Rutten and Ten Tusscher, 2021). We modified this auxin-PLT model to include the contribution of PLT auto-activation on PLT transcription. In this manuscript we present an auxin-PLT-JANUS-GIF1 model (baseline model) where PLT auto-activation is mediated through the PLT-JANUS protein complex (Equation 16). As a control (Figure S3), we also generated an auxin-PLT-autoactivation model in which unbound PLT positively regulates its own expression (Equation 17).

#### Auxin-PLT-JANUS-GIF1 model

In our baseline model, PLT production is positively regulated by two terms that together control a maximum production of PLT: a slow auxin regulation (Mähönen et al., 2014) and a rapid auto-activation through the PLT-JANUS complex:

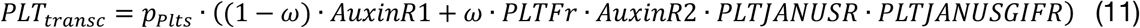

To keep PLT production constant, the contribution of each is defined by ω [0,1], such that the former is multiplied by 1-ω and the latter by, ω. Both fractions are constrained by auxin, based on previous modelling research showing that the auxin reflux loop constrains PLT responses (Rutten and Ten Tusscher, 2021; Figure S1B). Yet, while auxin regulation of PLT is a slow response, we assumed the PLT-JANUS regulation is a faster auxin response (Durgaprasad et al., 2019). The PLT-autoactivation term requires a fast auxin signalling input (B, Equation 4), otherwise the expression and distribution of PLT gradually expand to the whole meristem (Supplementary Figure S1B).

PLT auto-activation is regulated positively by PLT-JANUS, and negatively by PLT-JANUS-GIF1, and by very high levels of unbound PLT (see terms in Equation 11):

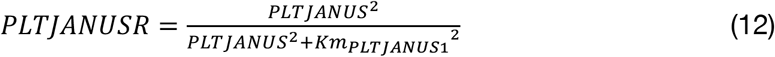

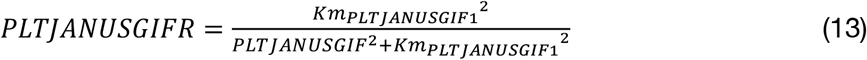

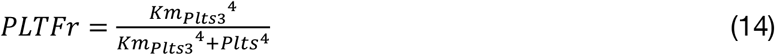

Importantly, we parameterize the model such that the repressive effects of unbound PLT are triggered only when it is found in extremely high levels (*Km*_*plts3*_), as those produced by inducible experiments (Durgaprasad et al., 2019). Indeed, a DEX term is included in PLT production to simulate the constitutive production of PLT upon induction, mimicking the DEX-inducible system, *35S::PLT2-GR* (Durgaprasad et al., 2019). Lastly, we included terms to describe the association and dissociation of PLT with TCP-SCR and JANUS:

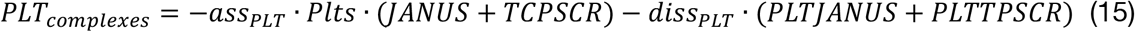

Together, this results in Equation 16, where we model DEX-induction of PLT, endogenous PLT transcriptional regulation, linear decay, and the association and dissociation of protein complexes:

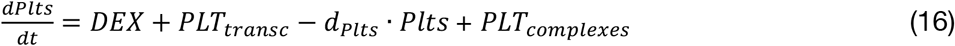

We used *ω*=0.6 as the default PLT auto-activation contribution accounting 60% of total PLT production (Figure S1C-F, and all DEX simulations). This value was defined based on the extent of the decrease in PLT2 expression in an inducible RNAi line targeting JANUS (Xiong et al., 2020).

Based on the size exclusion limitations of the plasmodesmata, we assumed that PLT cell-to-cell movement is only possible when it is in free form.

#### Simple auxin-PLT autoactivation model

The auxin-PLT model is a simplified version of the baseline model in which PLT auto-activation is mediated directly by unbound PLT, and no protein complexes are formed (Figure S3):

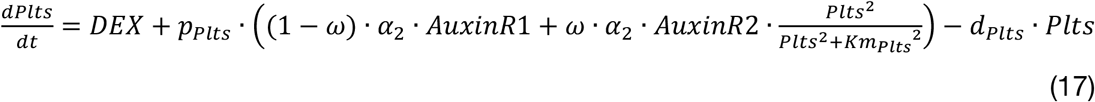

For this version of the model, we reduced the production of PLT to recover a constrained PLT gradient. This adjustment was necessary as this version of the model included no PLT protein complexes that limit the availability of PLT to move from cell to cell.

### JANUS, GIF1, TCP and SCR regulation

We included basal production of JANUS in all the simulated root cells to reproduce the homogenous expression of JANUS in the meristem (*Xiong et al., 2021*), and its linear decay. A DEX term was included for the simulation of constitutive production of JANUS upon induction, mimicking the inducible DEX line studied in this paper. To simulate the *janus* loss of function mutant we set the production of JANUS (*p*_*JANUS*_) to zero. We also include terms to describe its association and dissociation from PLT:

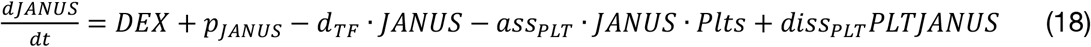

We defined basal expression of TCP20 and TCP21 (TCP hereafter) in all cells of the simulation (Shimotohno et al., 2018), and SCR expression only in the cell layer surrounding the central stele (Di Laurenzio et al., 1996; Shimotohno et al., 2018). In the case of SCR, we included a DEX term for constitutive production of SCR upon induction (Equation 20), mimicking the inducible DEX lines studied in this paper. Terms for linear decay and for their association and dissociation from the TCP-SCR complex were also included:

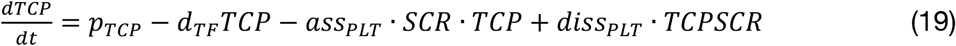

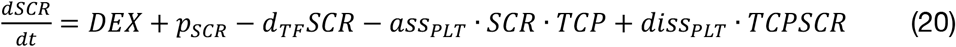

The TCP-SCR protein complex associates with PLT to form the PLT-TCP-SCR protein complex, and dissociates back to TCP-SCR.

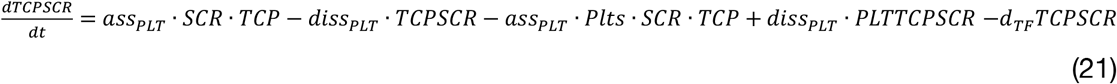

### PLT protein complexes

PLT forms protein complexes with different proteins in our model: PLT forms a complex with JANUS into the PLT-JANUS complex, GIF1 with the PLT-JANUS complex into the PLT-JANUS-GIF1 complex, and PLT with TCP-SCR to form the PLT-TCP-SCR protein complex. We assumed that the complexes inherit the decay rate of the most stable protein, which in this case is PLT:

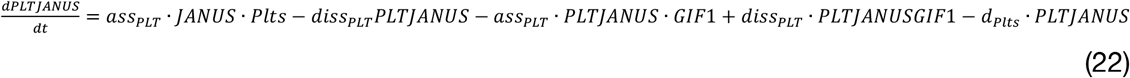

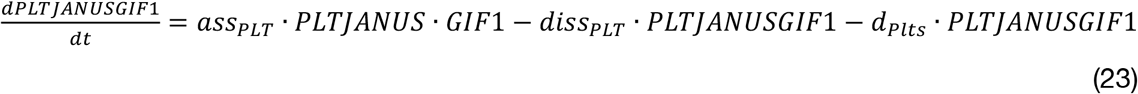

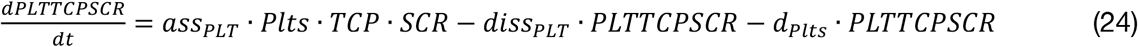

### PLT targets

In the Auxin-PLT-JANUS-GIF1 model we defined that the transcription of PLT targets is mediated by a PLT protein complex and repressed by unbound PLT. This was implemented such that the PLT-dependent production term in Equations 11 (PLT transcription), 25 (GIF1 production), 26 (WOX5 production), 28 (YUC3 production), is multiplied by *PLTFr* (Equation 14).

PLT regulates GIF1 expression, as evidenced by the abolishment in the signal of a GIF1 reporter in the *plt1 plt2* double mutant (Ercoli et al., 2018). Accordingly, the transcriptional regulator GIF1 is highly expressed in the stem cell niche (Ercoli et al., 2018). In our model, we included this PLT regulation through the PLT-JANUS protein complex, although other yet to be identified complex could be mediating this interaction. We also included a negative regulation by very large levels of free PLT (*PLTFr*) terms to account for GIF1 linear decay, and for its association and dissociation with the PLT-JANUS and PLT-JANUS-GIF1 complexes, respectively. The production of GIF1 (*p*_*GIF1*_) was set to zero for the simulation of *gif1* loss of function mutant.

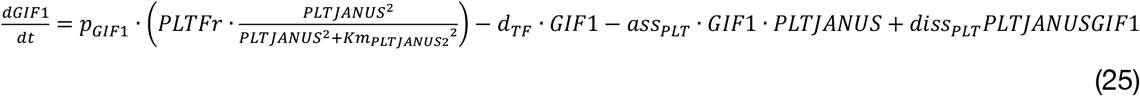

WOX5 production is positively regulated by the PLT-TCP-SCR protein complex, negatively by very large levels of free PLT (*PLTFr*), and it linearly decays.

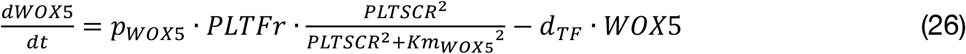

PLT-regulation of YUC3 is mediated in the model via the PLT-JANUS protein complex (although a yet to be discovered PLT-complex could be mediating this regulation).

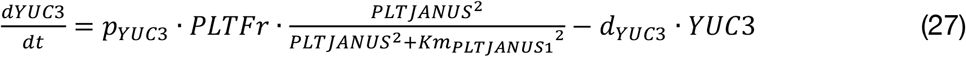

In the case of the simplified Auxin-PLT autoactivation model (Figure S3), this regulation is mediated directly by unbound PLT.

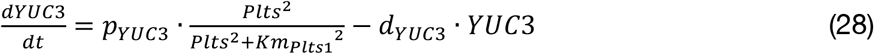

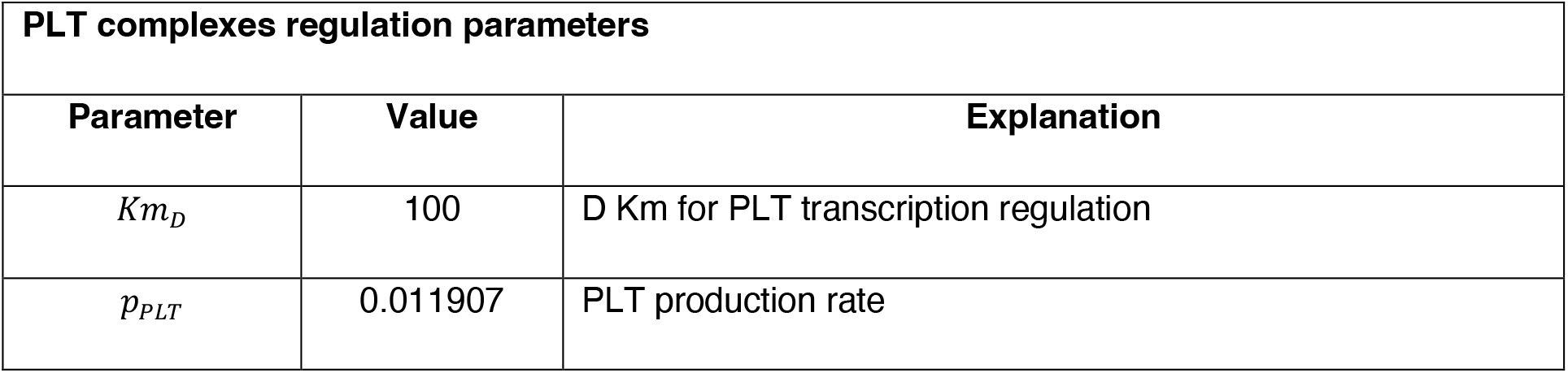
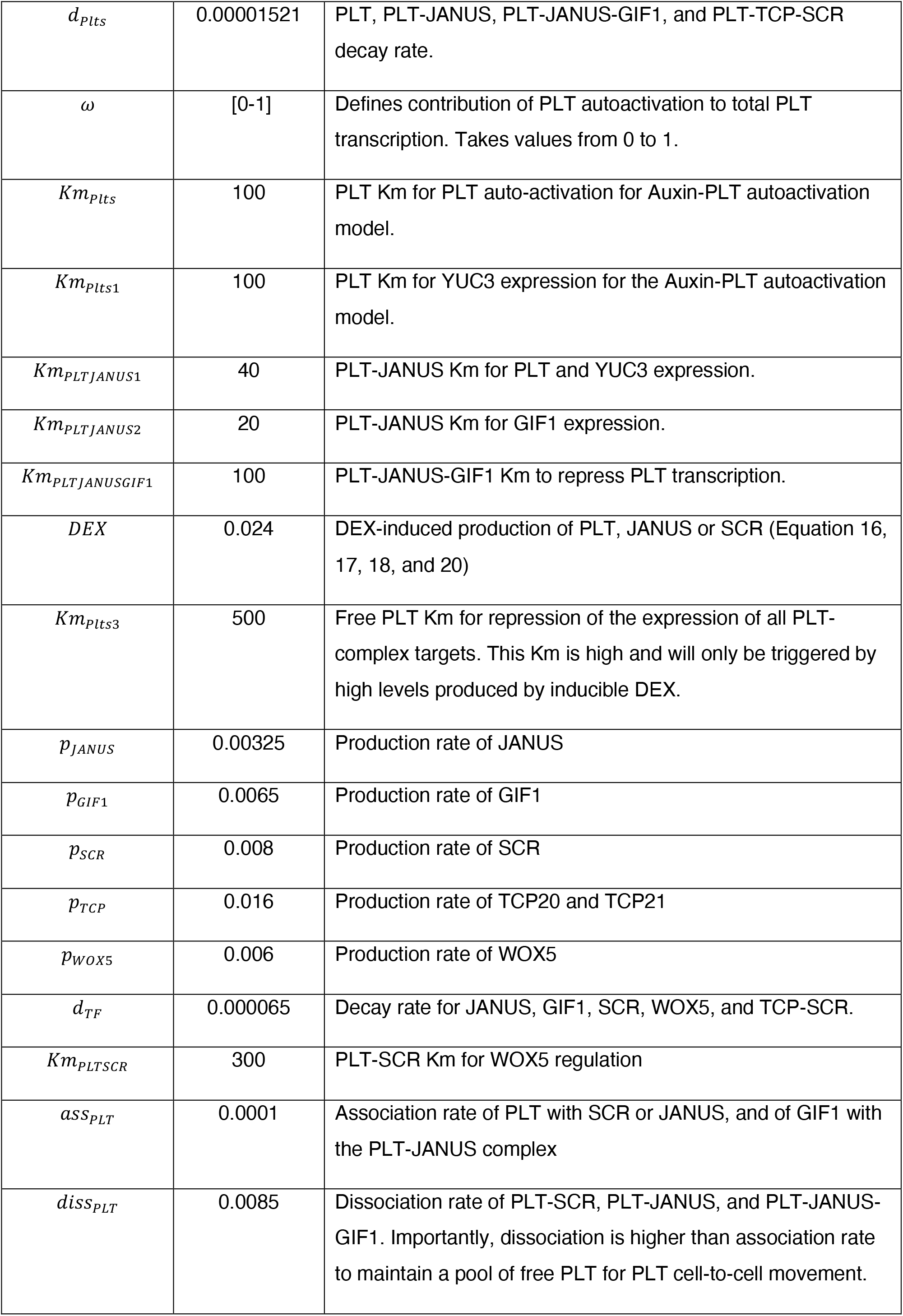

**PLT complexes regulation parameters**
| Parameter | Value | Explanation |
| --- | --- | --- |
| $Km_D$ | 100 | D Km for PLT transcription regulation |
| $p_{PLT}$ | 0.011907 | PLT production rate |
| $d_{PLTs}$ | 0.00001521 | PLT, PLT-JANUS, PLT-JANUS-GIF1, and PLT-TCP-SCR decay rate. |
| $\omega$ | [0-1] | Defines contribution of PLT autoactivation to total PLT transcription. Takes values from 0 to 1. |
| $Km_{PLTs}$ | 100 | PLT Km for PLT auto-activation for Auxin-PLT autoactivation model. |
| $Km_{PLTs1}$ | 100 | PLT Km for YUC3 expression for the Auxin-PLT autoactivation model. |
| $Km_{PLTJANUS1}$ | 40 | PLT-JANUS Km for PLT and YUC3 expression. |
| $Km_{PLTJANUS2}$ | 20 | PLT-JANUS Km for GIF1 expression. |
| $Km_{PLTJANUSGIF1}$ | 100 | PLT-JANUS-GIF1 Km to repress PLT transcription. |
| $DEX$ | 0.024 | DEX-induced production of PLT, JANUS or SCR (Equation 16, 17, 18, and 20) |
| $Km_{PLTs3}$ | 500 | Free PLT Km for repression of the expression of all PLT-complex targets. This Km is high and will only be triggered by high levels produced by inducible DEX. |
| $p_{JANUS}$ | 0.00325 | Production rate of JANUS |
| $p_{GIF1}$ | 0.0065 | Production rate of GIF1 |
| $p_{SCR}$ | 0.008 | Production rate of SCR |
| $p_{TCP}$ | 0.016 | Production rate of TCP20 and TCP21 |
| $p_{WOX5}$ | 0.006 | Production rate of WOX5 |
| $d_{TF}$ | 0.000065 | Decay rate for JANUS, GIF1, SCR, WOX5, and TCP-SCR. |
| $Km_{PLTSCR}$ | 300 | PLT-SCR Km for WOX5 regulation |
| $ass_{PLT}$ | 0.0001 | Association rate of PLT with SCR or JANUS, and of GIF1 with the PLT-JANUS complex |
| $diss_{PLT}$ | 0.0085 | Dissociation rate of PLT-SCR, PLT-JANUS, and PLT-JANUS-GIF1. Importantly, dissociation is higher than association rate to maintain a pool of free PLT for PLT cell-to-cell movement. |

### Regulation of auxin transporters

The PIN polarity pattern in each cell of the simulation is predefined and fixed based on their cell type and position along the simulated root apex. This is also the case for the influx transporters AUX1 and LAX3, while their levels are regulated by auxin:

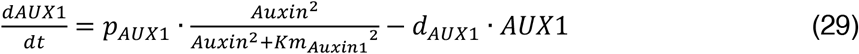

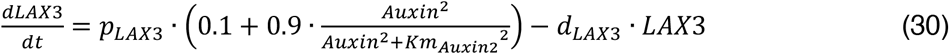

| Regulation of auxin transporters |  |  |
| --- | --- | --- |
| Parameter | Value | Explanation |
| $p_{YUC3}$ | 0.01 | YUC3 production rate |
| $d_{YUC3}$ | 0.0001 | YUC3 decay rate |
| $Km_{Auxin1}$ | 100 | Auxin Km for AUX1 regulation |
| $p_{AUX1}$ | 0.01 | AUX1 production rate |
| $d_{AUX1}$ | 0.0001 | AUX1 decay rate |
| $Km_{Auxin2}$ | 75 | Auxin Km for LAX3 regulation |
| $p_{LAX3}$ | 0.01 | LAX3 production rate |
| $d_{LAX3}$ | 0.0001 | LAX3 decay rate |
| Auxin transport and diffusion is simulated at the grid level as in our previous implementations of the model (Mähönen et al., 2014; Salvi et al., 2020). |  |  |
| Parameter | Value | Explanation |
| $D_{auxC}$ | 600 | Diffusion in cells. |
| $D_{auxW}$ | 40 | Diffusion in walls. |
| $eff$ | 1 | Passive auxin efflux |
| $InAux$ | 2.5 | Passive auxin influx |
| $dAux$ | 0.000052 | Auxin degradation rate |
| $pAux$ | 0.0005 | Basal auxin production rate |
| pPUMP | 0.2 | Active efflux rate |

### Root cut simulations

To simulate a root cut in our computer model we defined a cut position (above the QC or at the end of the meristem) and removed the cells below it. We closed the boundary of the distal cells to the simulated wound, to ensure they do not transport auxin or PLT to the medium. Consequently, both HC and LC resulted in the interruption of auxin reflux, and the accumulation of auxin at the wound site (Movie 2). Previous experimental observations showed that at 8 hpc a rapid upregulation of PLT2 transcription occurs in the central cells of the vasculature near the wounding site, and that this is an early marker of the regeneration competence of roots (Durgaprasad *et al*., *2019*). Thus, we ran our simulations for up to 8 hours after the cut (hpc) and defined the regeneration threshold as the level of PLT transcription in a vascular cell at the wound site following a LC (grey dashed lines in regeneration competence plots). Importantly, this timeframe is before significant growth deformation occurs, nor closure of plasmodesmata takes place (Mathew et al., 2025; Cohen et al., 2025). Thus, we do not model the actual restoration of the root tip shape as this occurs at a later point (*Matthews et al*., *2025*) To assess the individual role of either the slow auxin or of the fast PLT-JANUS in controlling the transcription of PLT in LC (Figure S1), we used ω=0.6, PLT-JANUS contribution to PLT transcription is 60%, as the baseline, and then set the role of either factor as zero after performing the cut.

## Authors contribution

M.L.G.G. and K.t.T. conceptualized the study. M.L.G.G. performed the computational simulations. M.L.G.G., K.P. and V.W. conceived the experiments. M.M., S.D. and C.R. performed the experiments. M.L.G.G. and K.t.T. wrote the original draft of the manuscript, K.P. edited, and all authors reviewed and approved the final version of the manuscript.

## Acknowledgements

This work was supported by the Netherlands Scientific Organization (Nederlandse Organisatie voor Wetenschappelijk Onderzoek, NWO) (ALWOP.542 to M.L.G.G and K.t.T). K.t.T. acknowledges VICI grant (VI.C.202.011 12268) (NWO). K.P. acknowledges the (BT/PR54722/BSA/33/509/2025) and (ANRF/ARG/2025/008174/LS). We are thankful to Jan Kees van Amerongen for his invaluable support with the management of computational facilities of the Theoretical Biology group (Utrecht University). We thank Maximilian W.D. Raas for useful discussions, and Lira van Dinther for inspiration for the title of this manuscript.

## Supplementary Figures

**Figure S1.**
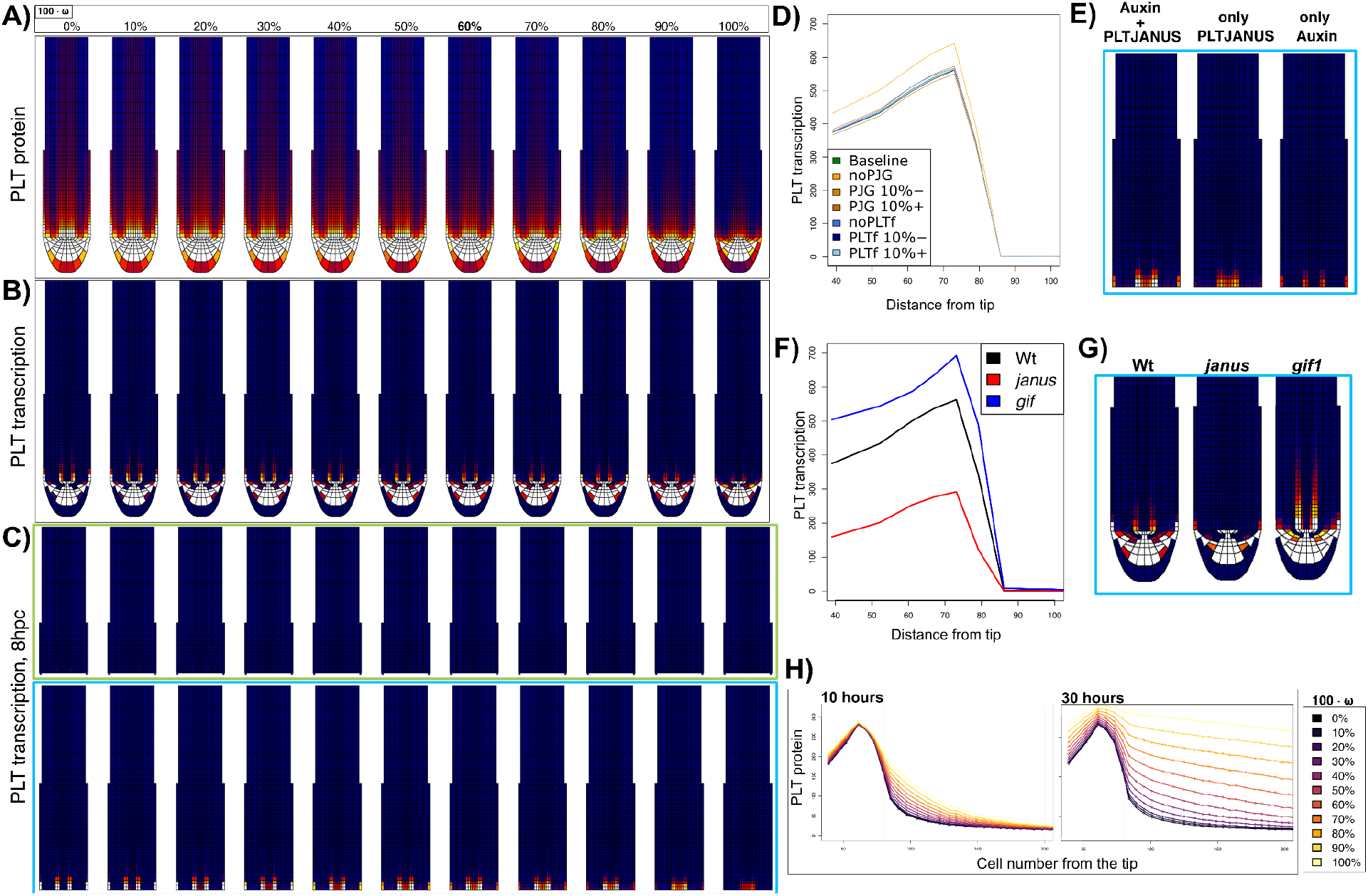
A) PLT protein, B) and PLT transcription produced by the Auxin-PLT-JANUS-GIF1 model with different contributions of PLT-JANUS over the regulation of *PLT* transcription (ω). C) PLT transcription 8 hours after a HC (green rectangle) or a LC (blue rectangle) was performed. Notice the *de novo* PLT transcription in the vascular cells of the wound requires PLT-JANUS regulation. (D) PLT transcription along a longitudinal file of vascular cells at the centre of the root for the baseline model, without PLT-JANUS-GIF1 repression of pPLT, with 10% less or more regulation of pPLT by PLT-JANUS-GIF1 (PJG 10%-, PJG 10%+), without the repressive effect of free PLT (noPLTf), and with 10% less or more regulation of PLT targets by free PLT (PLTf 10%-, PLTf 10%+). E) PLT transcription in the root apices 8 hours after a LC was performed for the baseline model (Auxin+PLTJANUS), or a version of the model where only PLT-JANUS or only auxin signalling regulates PLT transcription (only PLTJANUS and only Auxin, respectively). F) PLT transcription along a longitudinal file of vascular cells at the centre of the root for the baseline model (WT), the *janus* and the *gif1* loss of function mutants. G) Root apices showing PLT transcription of WT, *janus*, and *gif1*. Simulations of panels D to G used ω=0.6, that is the contribution of the PLT-JANUS of PLT transcription regulation is 60%. H) Protein levels along a longitudinal file of vascular cells at the centre of the root after 30 hours of simulation for different ω values, in a version of the model without fast auxin response constraining PLT auto-activation. Notice that when PLT auto-activation contribution is zero, and thus auxin contribution is 1, PLT is centred in the root stem cell niche and a graded PLT protein pattern is formed, as described previously (Mähönen et al., 2014; Salvi et al., 2020; Rutten and Ten Tusscher, 2021). As the fractional contribution of PLT is increased beyond marginal levels, PLT confinement is compromised and takes over the whole simulated root apex.

**Figure S2.**
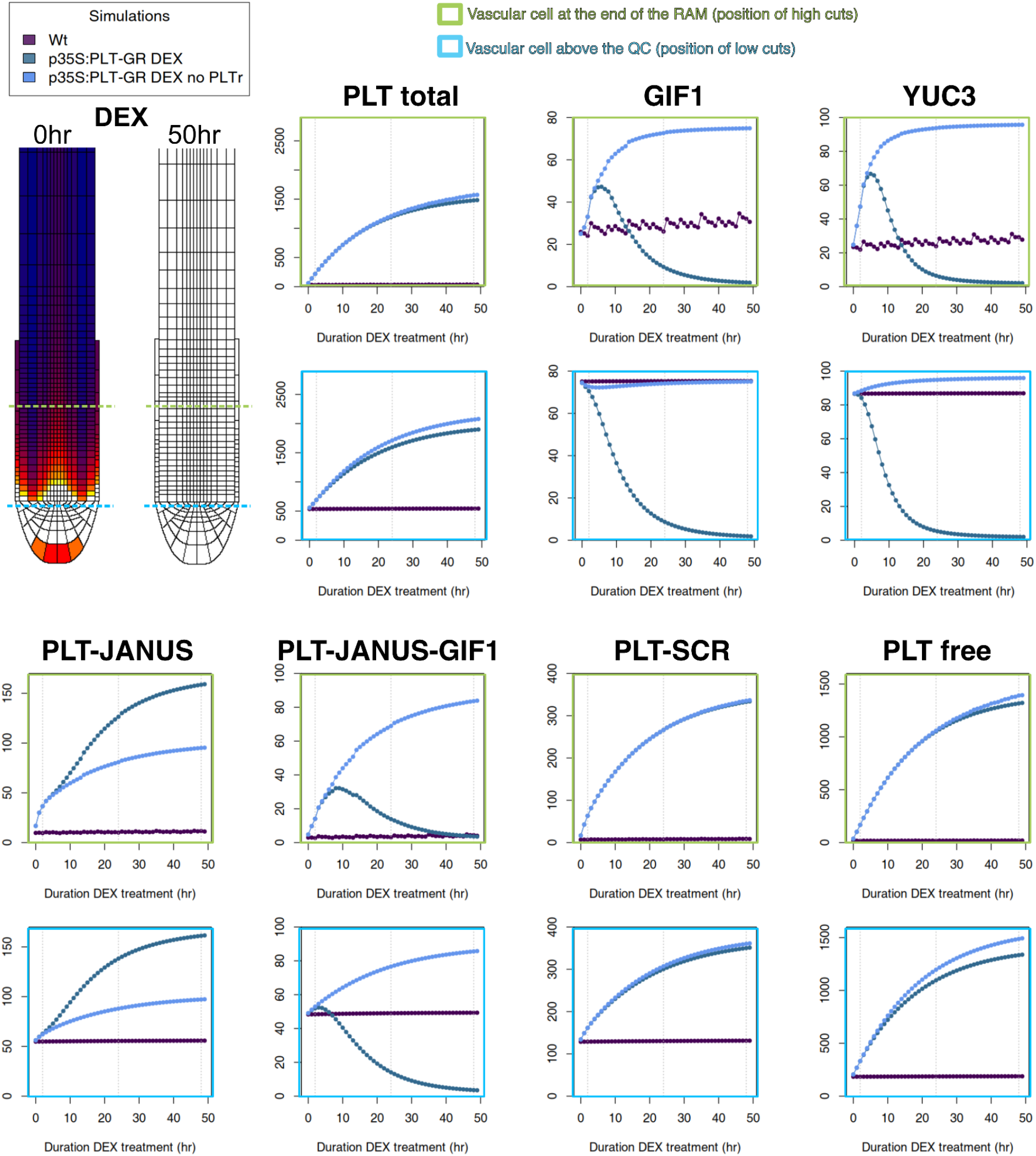
p35S::PLT-GR simulation. The levels of different elements of the regulatory network are plotted for two cells located at different positions within the meristem throughout 50 hours of DEX-induction for *p35S::PLT-GR* simulation. Green and blue lines in the roots mark the cells analysed, as this is the position of HC and LC are performed. Top: Total PLT protein levels, and expression of gene PLT targets (GIF1 and YUC3). Bottom: Levels of various PLT protein complexes, PLT-JANUS, PLT-JANUS-GIF1 and PLT-SCR (PLT-TCP-SCR), and of PLT in free form. Notice that despite the very high levels of PLT-JANUS in the *p35S::PLT-GR* simulation, the expression of various targets (e.g. GIF1 and YUC3) drops, and this response is lost in the *p35S::PLT-GR* without the repressive role of PLT in free form (‘no PLTr’). Grey vertical lines mark 0, 2h and 48h DEX.

**Figure S3.**
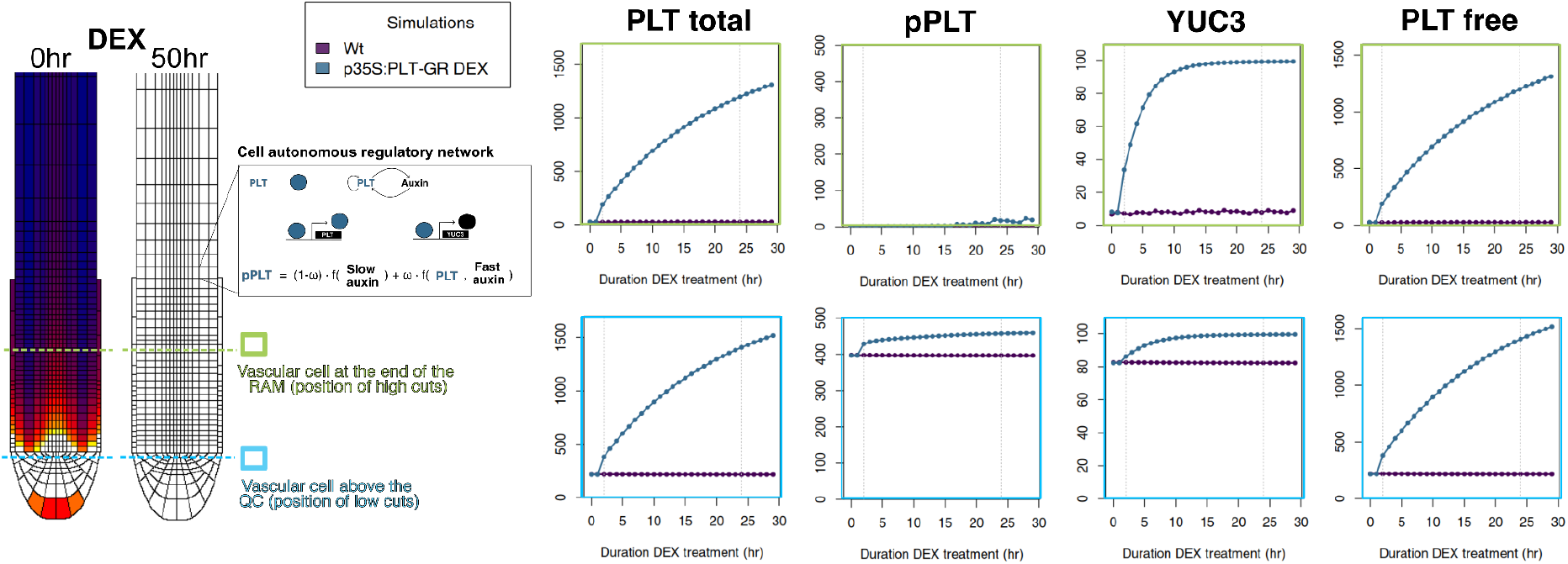
(A) Root model including PLT2 expression regulated slowly by auxin, and directly by PLT (auto-activation). The contribution of each is determined by ω. The levels of different elements of the regulatory network when simulating p35S::PLT-GR are plotted for two cells located at different positions within the meristem throughout 50 hours of DEX-induction. Green and blue lines in the roots mark the cells analysed. Top: Total PLT protein levels. Bottom: Levels of PLT expression, YUC3 and free PLT levels. Notice how this model exhibits monotonic responses to PLT dosage. Grey vertical lines mark 0, 2h and 48h DEX.

**Figure S4.**
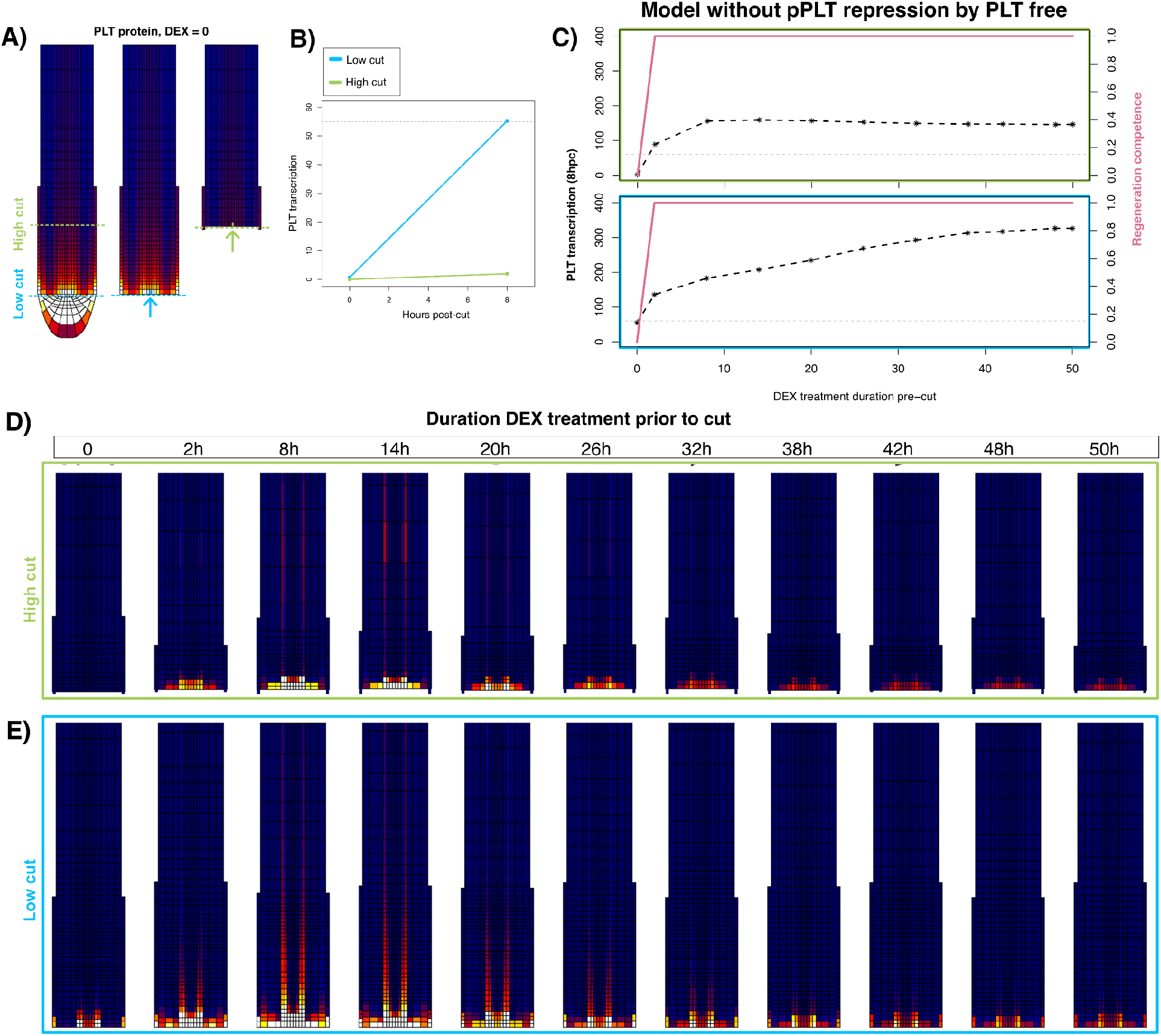
Effect of continuous induction of PLT by DEX on regeneration competence. (A) PLT protein levels in intact roots, and in roots after a LC or a HC. (B) PLT transcription right after a cut, and 8 hours later, for LC and HC. The value of PLT transcription reached in LC is used as the threshold needed for regeneration competence (dashed grey line). (C) Model without the negative role of free PLT on the regulation of PLT targets: PLT transcription in a vascular cell close to the wound 8 hours post-cut in roots treated with DEX for different durations, and their competence (threshold shown as a horizontal grey line). Notice the monotonic response in this model version. (D and E) PLT transcription in the root apices of HC and LC after 8hpc for different durations of DEX induction prior to the cut for the auxin-PLT-JANUS-GIF1 model.

**Figure S5.**
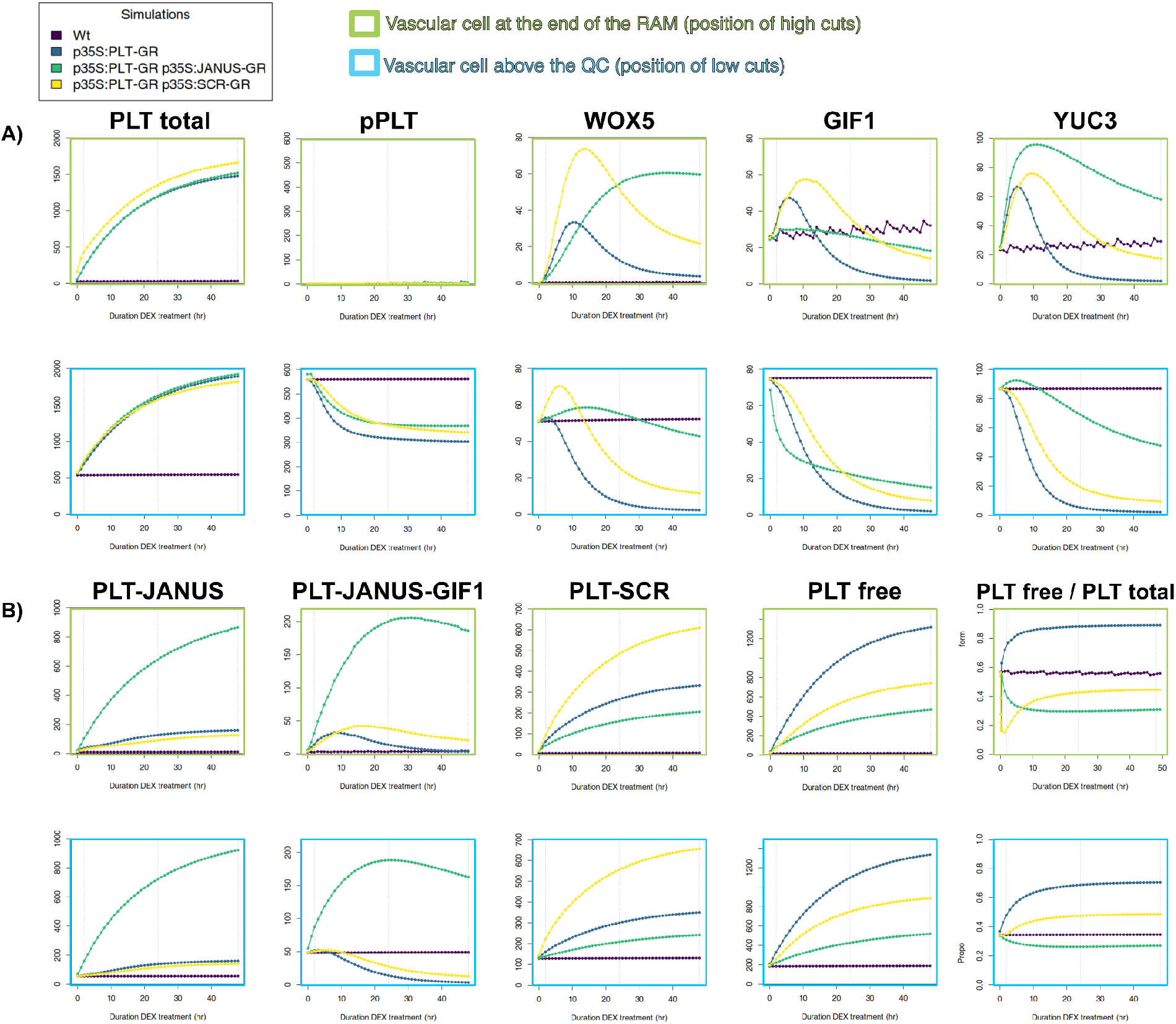
Effects of co-induction PLT alone or with an interacting protein. The levels of different elements of the regulatory network are plotted for two cells located at different positions within the meristem (light green marks a cell at the end of the meristem, and blue a cell above the QC) throughout 50 hours of DEX-induction for p35S::PLT-GR simulation alone or combined with either JANUS or SCR induction. Top: Total PLT protein levels, and expression of gene PLT targets (pPLT, WOX5, GIF1 and YUC3). Bottom: Levels of various PLT protein complexes, PLT-JANUS, PLT-JANUS-GIF1 and PLT-SCR (PLT-TCP-SCR), and of PLT in free form. The proportion of PLT free is also shown. Grey vertical lines mark 0, 2h and 48h DEX.

**Figure S6.**
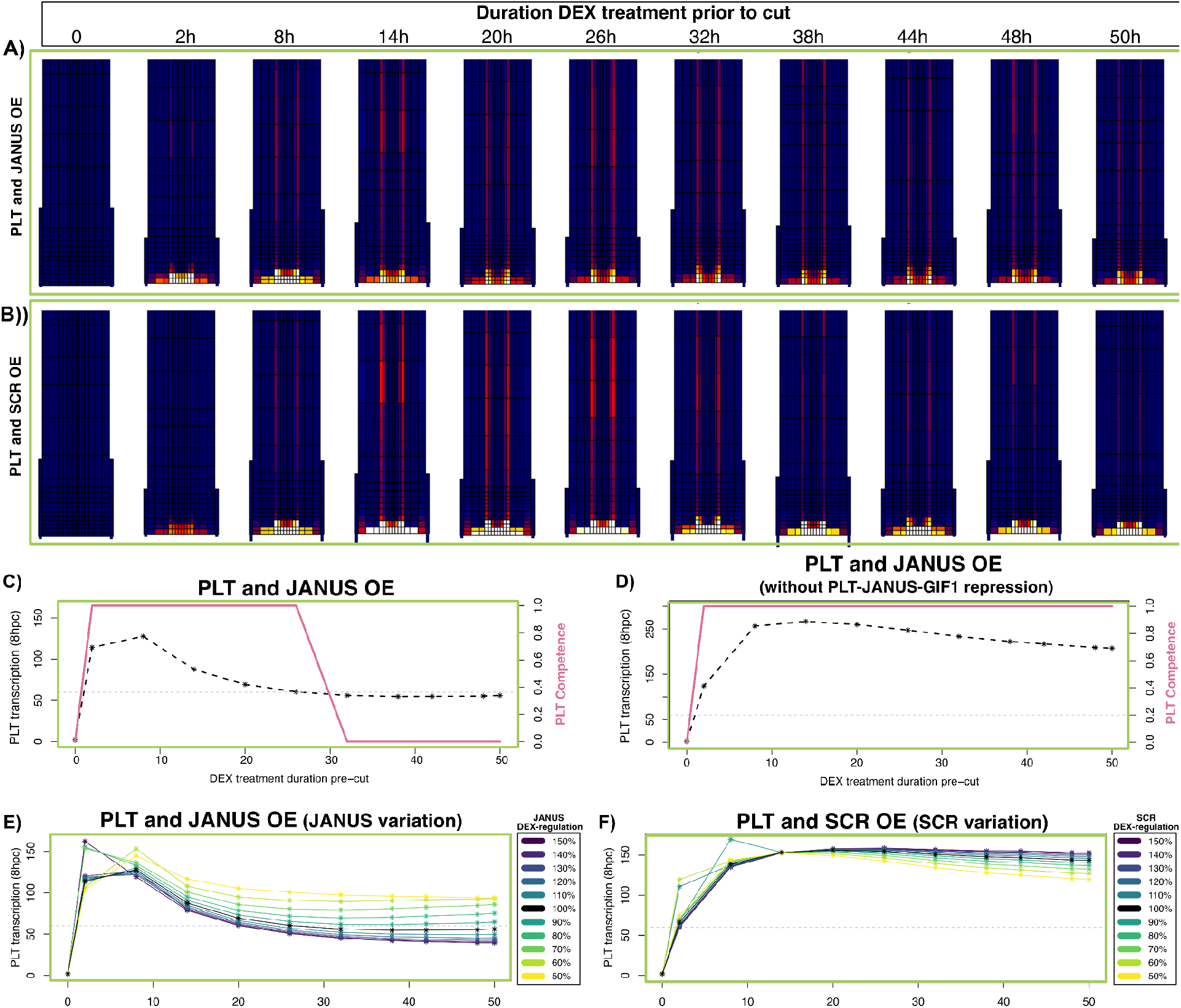
Cut simulations after co-induction of PLT with an interacting protein. PLT transcription in the root apices of HC after 8hpc for different durations of DEX induction prior to the cut for PLT and JANUS co-induction (A), and PLT and SCR co-induction (B) (Related to figure 4). (C) PLT transcription and regeneration competence of HC after the induced co-induction of PLT and JANUS for the baseline model, (D) and for a model version where PLT-JANUS-GIF1 does not repress PLT transcription. Effect of variation in the extent of induction of JANUS (E) and SCR (F) on regeneration competence of HC. The value for baseline simulations is 100%.

**Figure S7.**
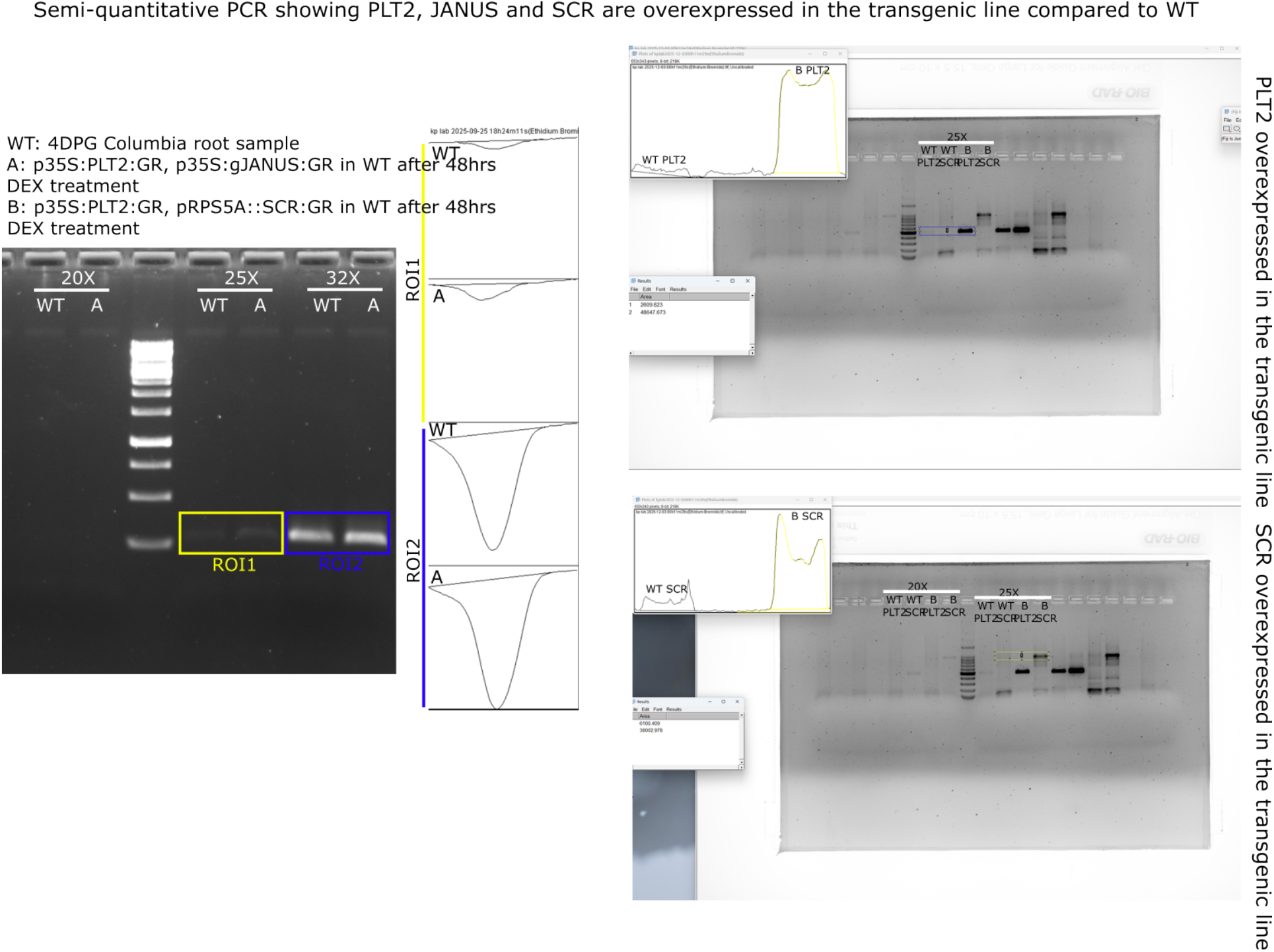
Quantification of PLT and JANUS or SCR expression in the p35S::PLT2-GR p35S::JANUS-GR, and p35S::PLT2-GR pRPS5a::SCR-GR transgenic lines.

## References

Aida, Mitsuhiro, et al. “The PLETHORA genes mediate patterning of the Arabidopsis root stem cell niche.” Cell 119.1 (2004): 109–120.

Baulies, Julia Lidia, et al. “MicroRNA control of stem cell reconstitution and growth in root regeneration.” Nature Plants 11.3 (2025): 531–542.

Birnbaum, Kenneth D., and Alejandro Sánchez Alvarado. “Slicing across kingdoms: regeneration in plants and animals.” Cell 132.4 (2008): 697–710.

Du, Yujuan, and Ben Scheres. “PLETHORA transcription factors orchestrate de novo organ patterning during Arabidopsis lateral root outgrowth.” Proceedings of the National Academy of Sciences 114.44 (2017): 11709–11714.

Durgaprasad, Kavya, et al. “Gradient expression of transcription factor imposes a boundary on organ regeneration potential in plants.” Cell Reports 29.2 (2019): 453–463.

Fallesen, T., et al. “Intermittent cell division dynamics in regenerating Arabidopsis roots reveals complex long-range interactions.” Quantitative Plant Biology 5 (2024): e7.

Galinha, Carla, et al. “PLETHORA proteins as dose-dependent master regulators of Arabidopsis root development.” Nature 449.7165 (2007): 1053–1057.

García-Gómez, Monica L., and Kirsten Ten Tusscher. “Multi-scale mechanisms driving root regeneration: From regeneration competence to tissue repatterning.” The Plant Journal 119.3 (2024): 1183–1196.

Heyman, Jefri, et al. “The heterodimeric transcription factor complex ERF115–PAT1 grants regeneration competence.” Nature plants 2.11 (2016): 16165.

Efroni, Idan, et al. “Root regeneration triggers an embryo-like sequence guided by hormonal interactions.” Cell 165.7 (2016): 1721–1733.

Ercoli María Florencia, et al. “GIF transcriptional coregulators control root meristem homeostasis.” The Plant Cell 30.2 (2018): 347–359.

Kareem, Abdul, et al. “PLETHORA genes control regeneration by a two-step mechanism.” Current Biology 25.8 (2015): 1017–1030.

Kerstens, Merijn, et al. “Two deeply conserved non-coding sequences control PLETHORA1/2 expression and coordinate embryo and root development.” Plant communications 6.10 (2025).

Kral, Nicolas, Alexandra Hanna Ougolnikova, and Giovanni Sena. “Externally imposed electric field enhances plant root tip regeneration.” Regeneration 3.3 (2016): 156–167.

Lee, Laura R., et al. “Glutathione accelerates the cell cycle and cellular reprogramming in plant regeneration.” Developmental Cell 60.8 (2025): 1153–1167.

Mähönen, Ari Pekka, et al. “PLETHORA gradient formation mechanism separates auxin responses.” Nature 515.7525 (2014): 125–129.

Mathew, Mabel Maria, and Kalika Prasad. “Model systems for regeneration: Arabidopsis.” Development 148.6 (2021): dev195347.

Mathew, Mabel Maria, et al. “Wound repair in plants guided by cell geometry.” Current Biology 35.16 (2025): 3851–3868.

Matosevich, Rotem, et al. “Local auxin biosynthesis is required for root regeneration after wounding.” Nature Plants 6.8 (2020): 1020–1030.

Omary, Moutasem, Rotem Matosevich, and Idan Efroni. “Systemic control of plant regeneration and wound repair.” New Phytologist 237.2 (2023): 408–413.

Rahni, Ramin, et al. “Histone deacetylases and cell-cycle regulators orchestrate cell-identity transitions during Arabidopsis root regeneration.” Molecular plant 19.5 (2026): 1036–1058.

Rodriguez, Ramiro E., et al. “MicroRNA miR396 regulates the switch between stem cells and transit-amplifying cells in Arabidopsis roots.” The Plant Cell 27.12 (2015): 3354–3366.

Rutten, Jacob P., and Kirsten H. Ten Tusscher. “Bootstrapping and pinning down the root meristem; the auxin–PLT–ARR network unites robustness and sensitivity in meristem growth control.” International Journal of Molecular Sciences 22.9 (2021): 4731.

Salvi, Elena, et al. “A self-organized PLT/Auxin/ARR-B network controls the dynamics of root zonation development in Arabidopsis thaliana.” Developmental cell 53.4 (2020): 431–443.

Santuari, Luca, et al. “The PLETHORA gene regulatory network guides growth and cell differentiation in Arabidopsis roots.” The Plant Cell 28.12 (2016): 2937–2951.

Sena, Giovanni, et al. “Organ regeneration does not require a functional stem cell niche in plants.” Nature 457.7233 (2009): 1150–1153.

Shimotohno, Akie, et al. “Root stem cell niche organizer specification by molecular convergence of PLETHORA and SCARECROW transcription factor modules.” Genes & development 32. 15-16 (2018): 1085–1100.

Strotmann, Vivien I., Monica L. García-Gómez, and Yvonne Stahl. “Root stem cell homeostasis in Arabidopsis involves cell-type specific transcription factor complexes.” EMBO reports 26.9 (2025): 2323–2346.

Xiong, Feng, et al. “Arabidopsis JANUS regulates embryonic pattern formation through Pol II-mediated transcription of WOX2 and PIN7.” Iscience 19 (2019): 1179–1188.

Xiong, Feng, et al. “Transcriptional regulation of PLETHORA1 in the root meristem through an importin and its two antagonistic cargos.” The Plant Cell 32.12 (2020): 3812–3824.

Xu, Jian, et al. “A molecular framework for plant regeneration.” Science 311.5759 (2006): 385–388.

